# Leveraging short and long-read sequencing to comprehensively characterize alternatively polyadenylated and alternatively spliced transcripts

**DOI:** 10.64898/2026.11.03.763780

**Authors:** Mahesh Gupta, Lael Pasipamire, Natoya Peart, Jie Li, Demiah Lockett, Isabella Todd, Haavia Rajakumar, Chioniso Patience Masamha

## Abstract

Pre-mRNA processing through alternative splicing and alternative polyadenylation generates diverse transcripts from the same gene. Alternative splicing involves use of different exons and specific 3’Untranslated regions (3’UTRs). When limited to the same terminal exon, 3’end formation through alternative polyadenylation gives rise to transcripts containing 3’UTRs of different sizes which affect their subcellular localization, stability, and translatability. The full repertoire of transcripts arising from alternative pre-mRNA processing under different biological and disease states remains unknown. Using the blood cancer, mantle cell lymphoma’s (MCL) as our experimental model we found that two genes, *SOX11* and *ATM,* that contribute to the disease pathology, express transcripts that have short 3’UTRs. Apart from select transcripts, our analysis of short-read sequencing data did not uncover widespread global shortening of 3’UTRs in MCL. However, we discovered that both the canonical and non-canonical polyadenylation signals (PASs), were used for 3’end formation. We experimentally verified the PASs used by specific transcripts. Combining short-read and long-read sequencing data enabled us to characterize, classify and identify alternatively splicing in MCL. We identified splice variants for several genes including *PTEN*, *EZH2* and validated the expression of GLS1 at the protein level and by using *GLS1* inhibitors. Collectively, our approach provide insights into how to identify alternatively spliced and alternatively polyadenylated transcripts.

## INTRODUCTION

Use of alternative promoters during transcription, as well as processing of pre-mRNAs through alternative splicing and alternative polyadenylation results in multiple transcript isoforms being formed from a single gene (1–3). During pre-mRNA processing, the spliceosome catalyzes various forms of alternative splicing including use of alternative 5’ donor splice sites, alternative 3’acceptor splice sites, mutually exclusive exons, intron retention and exon skipping (cassette exon) (3,4). Alternative splicing results in multiple protein products arising from a single gene and about 95% of mammalian genes are multi-exon (3,5). Dysregulation of alternative splicing is associated with several diseases including cancer (as reviewed (6,7). Besides splicing, another related co-transcriptional process, 3’end formation, is also involved in pre-mRNA processing. This involves 3’end cleavage downstream of polyadenylation signal (PAS), usually A(A/U)UAAA, followed by polyadenylation culminating in the addition of a poly(A) tail to the transcript (8). Usage of alternative PASs through alternative polyadenylation (APA) further adds to transcript diversity by generating transcripts with different 3’ends. Up to 70% of human genes contain multiple PASs (9–11). When APA occurs, it may involve splicing, use of intronic PASs or alternative PASs within the same terminal exon (12). The later type of APA (3’UTR-APA) which does not affect the open reading frame is the most common (13). At first look 3’UTR-APA appears innocuous until you focus on the makeup of the 3’UTR. The 3’UTR contains sequences that allow recruitment and binding of RNA binding proteins as well as miRNAs and shortening the 3’UTR eliminates some of these regulatory sequences (14,15). As a result, altering the size of the 3’UTR may alter the transcripts translatability and its stability, subsequently impacting the protein dosage, as well as its subcellular localization and thus affecting its spatial expression and function (16–20). Dysregulation of APA contributes to different pathological states including cancer (as reviewed (21,22)). Identification of transcripts arising from alternative splicing and alternative polyadenylation in different cell types and cell states, as well as their contributions to different biological and pathological states is still ongoing.

For this project, we used the B-cell hematological malignancy, Mantle Cell Lymphoma (MCL) as our experimental model to investigate alternative splicing and alternative polyadenylation due to its genomic complexity. MCL is divided into various subcategories with different clinical presentations and molecular profiles (23,24). The initiating lesion in the majority of MCL (∼95%) is a t(11;14) (q14;q32) translocation event that constitutively activates the expression of the initiating oncogenic cell cycle regulatory protein cyclin D1 (*CCND1*) expression in mature peripheral B-cells (25). After this transactivation event, a variety of *CCND1* transcripts are formed in MCL due to alternative splicing and alternative polyadenylation which affects protein expression. The *CCND1* gene undergoes alternative splicing to generate the two isoforms (26). One of them is the 1.7kb cyclin D1b isoform which lacks exon 5 and retains a section of intron 4. The second is the 4.5kb-cyclin D1a isoform which results from normal splicing of all 5 exons (27). Since the cyclin D1b protein is rarely detected in MCL and is not regarded as important for MCL pathogenesis, the isoform being referred to as *CCND1*(cyclin D1) in this manuscript will always be the cyclin D1a form (*CCND1*) (28,29). The oncogenic *CCND1* splice isoform also undergoes 3’UTR-APA in MCL through diverse mechanisms to give rise to *CCND1* transcripts with truncated 3’UTR which are associated with increased *CCND1* stability, increased protein expression, hyperproliferative disease and a reduction in patient survival (30,31). Besides *CCND1*, not much is known about alternative pre-mRNA processing in MCL. A major focus has been the identification of mutations in driver genes that contribute to MCL pathogenesis as well as chromosomal deletions involving tumor suppressors (32–35). Recurrent mutations in several genes including *CCND1*, *ATM, BTK, ARID2, CARD11, CDKN2A, CHD2, HNRNPH1, KMT2D, MAP3K14, NOTCH1/2, NSD2, RIMS2, SMARCA4, SP140, SPEN, S1PR1, TP53, TRFAF2, TLR2*, are associated with MCL oncogenesis (32–35). Apart from mutations, changes in the expression levels of several genes including *SOX11*, *MYC* (c-myc), and *TP53* have been reported in MCL (36–39).

In terms of splicing, recurrent mutations in the RNA binding protein *HNRNPH1* resulting in intronic splicing were associated with an increase in protein levels of the protein in MCL patient tissues (40).

The goal of this paper was to look at the MCL transcriptome by looking at transcripts resulting from alternative polyadenylation and alternative splicing. In this study, we compared gene expression in different human MCL cell lines using short-read RNA Sequencing (RNA-Seq). We determined the global usage of different polyadenylation signals (PASs) and identified the specific PASs used for select genes using 3’RACE. Comparisons of APA within the same terminal exon (3’UTR-APA) was done using the percentage change of distal poly(A) site usage index (PDUI) (41,42) for the mantle cell lymphoma cell line RNA-Seq data used in the study as well as publicly available RNA-Seq data for B-cells from normal individuals. To fully characterize the transcripts arising from alternative splicing we integrated our RNA-Seq. data with our long-read Iso Sequencing data (Iso-Seq) for select genes including *GLS1*, a critical driver of cancer metabolism. We used qRT-PCR and Western blot analysis to further interrogate expression of alternatively spliced the glutaminase (*GLS1*) transcripts and the impact of using glutaminase inhibitors on the growth of MCL cell lines. Taken together, our data identified transcripts arising from alternative pre-mRNA processing in MCL and our approach can be used to characterize other transcriptomes.

## MATERIAL AND METHODS

### Cell culture and cell viability assays

Human primary CD19+ B-cells and the mantle cell lymphoma cell lines Jeko-1 (Catalog# CRL-3006), Rec-1 (Catalog# CRL-3004), Mino (Catalog# CRL-3000), Z-138 (Catalog# CRL-3001) and the B-lymphocyte cell line RPMI-1788 (Catalog# CCL-156) B-cell line were purchased from ATCC (Manassas, Virginia). The Granta-519 mantle cell lymphoma cell line was purchased from DSMZ (The Leibniz Institute DSMZ, Braunschweig, Germany, Catalog# ACC 342). The B-cells and the RPMI-1788 cell line were cultured in RPMI-1640 cell culture media (ATCC, Manassas, Virginia, Catalog#30-2001) while all the mantle cell lymphoma cell lines were cultured in DMEM GlutaMAX™ cell culture medium (Thermo Fisher Scientific, Waltham, MA, Catalog#10569044). Both types of cell culture media were supplemented with 10% heat-inactivated FBS and 5% Penicillin-Streptomycin (Catalog#15140122) from Thermo Fisher Scientific (Waltham, MA). Human Induced Pluripotent Cells, hIPSCs, and human peripheral B-cells were purchased from StemCell Technologies (Vancouver, BC, Catalog#200-0769 and 70023). The hIPSCs were maintained in mSTeR Plus Medium from StemCell Technologies (Vancouver, BC, Catalog #100-0276) and the B-cells were grown and maintained in ImmunoCult Human B Cell Expansion Kit from StemCell Technologies (Vancouver, BC, Catalog#100-0645). Cells were grown at 37°C in a 5% CO_2_ humidified incubator. For cell viability assays, an equal number of cells were plated in 96-well plates (10,000 cells/plate). After 24 hours, cells were treated with DMSO or different concentrations of glutaminase inhibitors for 48 hours. Cell viability was determined using the MTT Assay (ATCC, Manassas, Virginia, Catalog#30-1010K).

### RNA Extraction and cDNA Synthesis

After TRIzol (Thermo Fisher Scientific, Waltham, MA, Catalog#15596018) extraction, total RNA was quantified and the 260/280 value determined using a Nanodrop. The total RNA was run on an ethidium-bromide-stained agarose gel to check RNA integrity. High quality RNA was then processed further as needed.

For qRT-PCR of SOX11, RNA was reverse transcribed using HiScript® III RT SuperMix for qPCR (Gene Biosystems, Burlington, ON, Catalog# R323-01) or HiScript® IV RT Supermix for qPCR (GeneBiosystems, Burlington, ON, Catalog# R423-01). Quantitative real time PCR was performed using Luna Universal qPCR Master Mix (NEB, Whitby, ON, Catalog#M3003S). Fold change of the distal UTR usage was calculated using a modified ΔΔCT. The CT values of the Distal and Common UTR were normalized to the CT of GAPDH to obtain ΔCT. The ΔCT obtained for the Distal and Common primers were adjusted using the calculated primer efficiencies. Primer efficiencies for SOX11 primers were assessed using genomic DNA from HCT116 cells. Then the ΔΔCT^test^ ^cell^ ^line^ was calculated by subtracting the Adjusted ΔCT_Common_ from the AdjustedΔCT_Distal_. The ΔΔΔCT was calculated by subtracting the ΔΔCT^B-cells/RPMI1788^ from ΔΔCT^test^ ^cell^ ^ine^, and the fold change with respect to B-cells (RPMI 1788) was calculated using 2^ΔΔΔCT^.

For *SOX11* 3’RACE, total RNA was treated with DNase I (NEB, Whitby, ON, Catalog#M0303S) according to manufacturer’s specifications. Following DNase treatment, the RNA was cleaned up with NEB’s Monarch RNA Cleanup Kits (Company, Whitby, ON, Catalog#T2030S). Following DNase treatment, 50% of the eluted RNA was reverse transcribed using MMLV Reverse Transcriptase (Gene Biosystems, Burlington, ON, Catalog# R1042) primer OligodT T7 5′-GGCCGGTAATACGACTCACTATAGTTTTTTTTTTTTTTTTTTTTTTTTT-3′. The remaining eluate was used for the negative control with no reverse transcriptase included.

To detect all *SOX11* 3’UTR isoforms, we used nested primers within the open reading frame and near the stop codon. To detect longer UTR isoforms of *SOX11*, we designed primers near the end of the within 200-500nts of the end of the RefSeq transcript for *SOX11* (NM_003108). The 3′RACE reactions were performed using Pfu polymerase (Gene Biosystems, Burlington, ON, Catalog#P1023) as previously described. Products of the 1st and 2nd PCR were resolved on 1% agarose gel in 1X TAE. To determine the cleavage products, the PCR products of the 2nd PCR were cloned with 5min TA/Blunt-Zero Cloning Kit (Gene Biosystems, Burlington, ON, Catalog#C601-01CC) according to manufacturer’s recommendations and positive clones were submitted for Sanger Sequencing at The Center for Applied Genomics at Hospital of Sick Children, Toronto, Canada.

For *ATM*, *BTK* and *BAX*, cDNA was reverse transcribed using RevertAid RT Reverse Transcription Kit (Thermo Fisher Scientific, Waltham, MA, Catalog#K1621). Random hexamer primers were used for cDNA used for PCR and qRT-PCR and the oligo(dT)_18_ primer was used to synthesize cDNA used for 3’ RACE and 3’ RACE was performed as previously described with slight modifications (31). In brief, for the first round of PCR, the transcript specific forward primer was used with oligo(dT_25_)T7primer and for the second round of PCR a nested transcript specific forward primer was used together with a T7 primer. However, Phusion high-fidelity DNA polymerase (Phusion High-Fidelity PCR Master Mix, Thermo Fisher Scientific, Waltham, MA, Catalog# F531L) was used instead of pfu polymerase. After gel purification of the second round PCR product using Wizard® SV Gel and PCR Clean-Up System (Promega, Madison, WI, Catalog# A9284), TOPO cloning was done using Zero Blunt TOPO PCR Cloning Kit for Sequencing (Thermo Fisher Scientific, Waltham, MA, Catalog# K280020). The resulting cloning products (and in some cases the gel purified PCR products) were sequenced using Sanger Sequencing (Lonestar Labs, TX).

### Library preparation and poly(A) RNA sequencing (RNA-Seq)

After in house quality control, high quality RNA samples were submitted to LCSciences (Houston, Texas site: https://lcsciences.com/services/rna-sequencing-services/rna-seq/) for Poly(A) RNA sequencing (RNA-Seq) and bioinformatic analysis. In brief, the integrity of the RNA samples was further analyzed using the Agilent Technologies 2100 Bioanalyzer. The polyadenylated transcripts containing a poly(A) tail were purified using oligo-(dT) magnetic, fragmented and the RNA-Seq libraries were prepared according to the Illumina’s TruSeq-stranded-mRNA sample preparation protocol. Quantification and quality control of the RNA-Seq library was performed using the Agilent Technologies 2100 Bioanalyzer High Sensitivity DNA Chip. This was followed by paired-ended sequencing using Illumina’s NovaSeq 6000 sequencing system.

After sequencing, the data was analyzed using LCSciences’ (Houston, Texas, https://lcsciences.com/services/rna-sequencing-services/rna-seq/) in-house optimized bioinformatic pipeline. In summary, after processing with Cutadapt (1.10) (43), read quality was verified using FastQC (v 0.10.1) (http://www.bioinformatics.babraham.ac.uk/projects/fastqc/). Reads were mapped to the human genome GRCh38 Ensembl 107 (v107) (ftp://ftp.ensembl.org/pub/release-107/fasta/homo_sapiens/dna/) using HISAT2 (v2.0) (44) and the mapped reads were assembled using StringTie (v.1.3.4) (45). Different expression analysis: StringTie was used determine mRNA expression levels by calculating FPKM and differential expression of mRNAs was performed using R package DESeq2 (46)/ by R package edgeR(47). Only transcripts with a false discovery rate (FDR) below 0.05 and absolute fold change ≥ 2 were considered differentially expressed. Pathway and process enrichment analysis was performed using Gene Orthology (GO) (http://geneontology.org).

### RNA-Seq poly(A) analysis

To investigate alternative polyadenylation for the short-read RNA-Seq data, quality control was performed to remove adapters and low quality bases using trim galore (v0.6.7). Alignment was done to human genome (GRCh38) using STAR(v2.7.10b) and bedgraph was created using bedtools (v2.29.2). Alternative polyadenylation analysis was carried out using APA analysis using DaPars2 (https://leilisysbio.github.io/DaPars2_Documentation/DaPars2.html).

### Iso-Sequencing data

Sample preparation and sequencing of long-read Iso-Seq data was previously described (48). Data is available in the ArrayExpress (BioStudies.) database (http://www.ebi.ac.uk/arrayexpress), accession number E-MTAB-17128.

### Integrating RNA-Seq and Iso-Seq data

R package rtracklayer (49) was used to combine the RNA-Seq coverage and the gene models generated using SQUANTI3 (50) on Iso-Seq data.

### PCR and qRT-PCR

Multiple forward and reverse primers were designed to target specific locations on transcripts downloaded from ensemble genome browser. After initial primer screening using Hot Start Taq 2X Master Mix (New England Biolabs, Ipswich, MA, Catalog# M0496L) select primers giving only one PCR product were chosen from each target region. Validation of primers was done using triplicate biological samples of Jeko-1 and Granta-519 mRNA. qRT-PCR was done using KAPA SYBR FAST qPCR Master Mix (2X) (Roche, Indianapolis, IN, Catalog# 7959397001) using at least 3 biological replicates. The sequences of primers targeting the open reading frames of the GAPDH (51), GAC and KGA (52), as well as nested reverse primers for 3’RACE (31) were previously described. The primers used for quantifying total *SOX11* transcripts were previously described (53). The rest of the primer sequences are provided as follows.

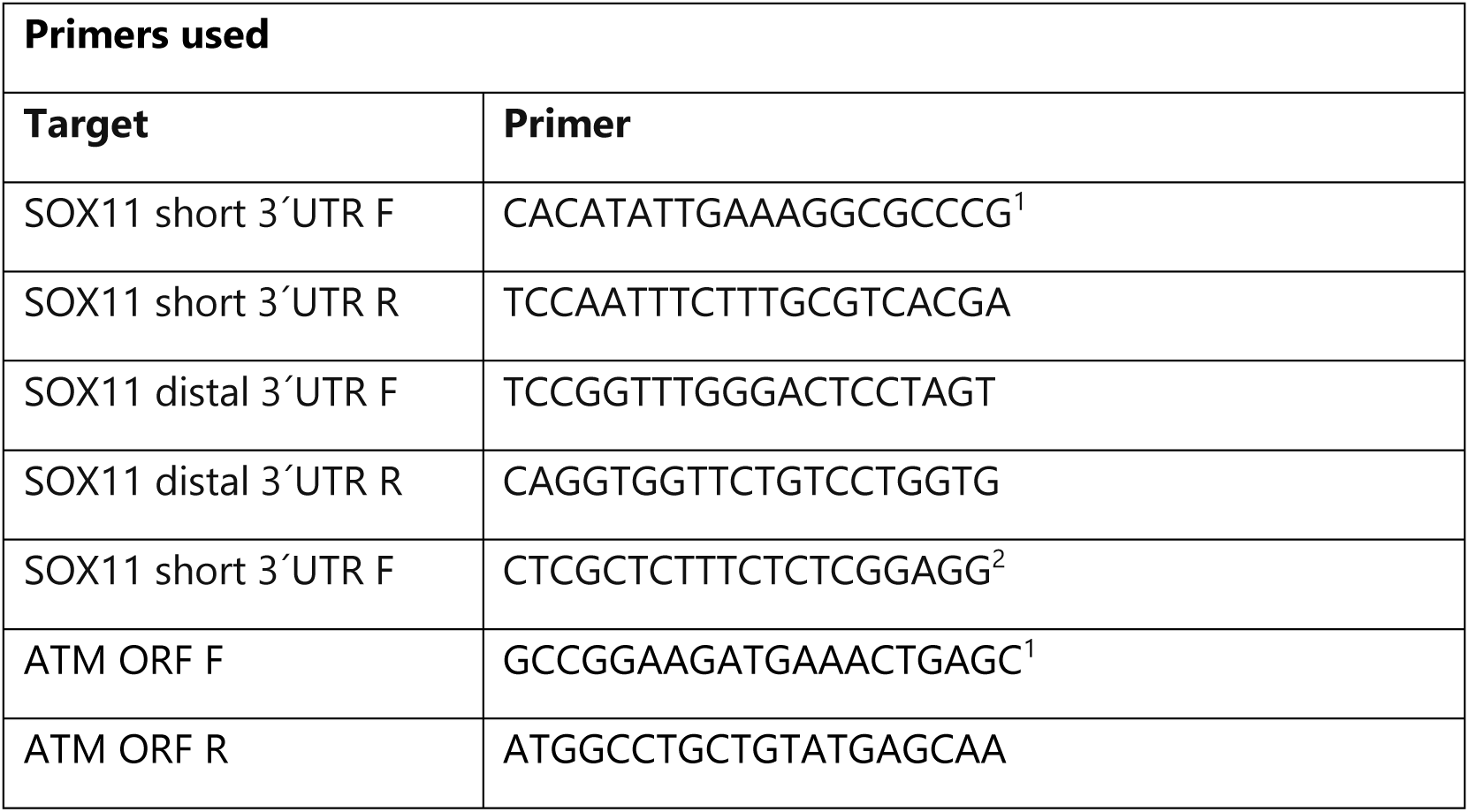
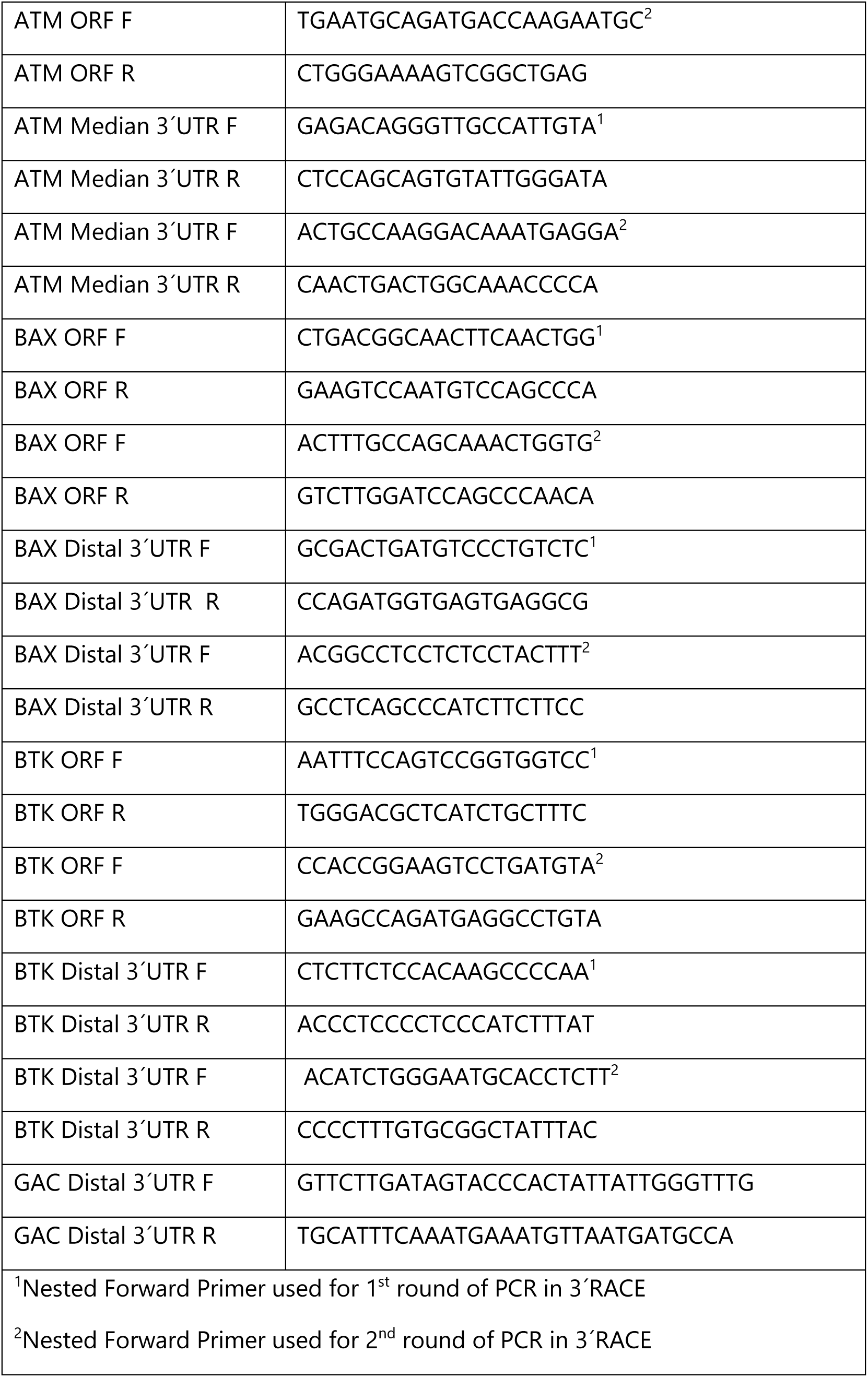

### Western blotting

Protein concentrations from cell lysates were quantified using Pierce™ Detergent Compatible Bradford Assay Kit (Thermo Fisher Scientific, Waltham, MA, Catalog#23246) and separated using SDS-PAGE. Proteins were transferred to a PVDF membrane (Bio-Rad, Hercules, CA, Catalog#1620177) and the membrane was blocked using 5% milk followed by incubation with anti-glutaminase (Abcam, Waltham, MA,Catalog# ab156876) and anti-GAPDH (Abcam, Waltham, MA, Catalog#ab128915). Blots were then washed and incubated with secondary Alexa Fluor™ 680 antibody (Thermo Fisher Scientific, Waltham, MA, Catalog# A27042). Fluorescence was detected using a Bio-Rad ChemiDoc Imaging System (Bio-Rad, Hercules, CA).

### Statistical analysis

GraphPad Prism was used for statistical analysis.

## RESULTS

### Differential gene expression

The Mantle Cell Lymphoma (MCL) subtype of non-Hodgkins Lymphoma is clinically considered a highly heterogenous disease (34) (and as reviewed (33,54)). In order to determine differences in gene expression in MCL, short-read RNA Sequencing (RNA-Seq) was carried out on three distinct human MCL cell lines, Jeko-1 (MCL1), Granta-519 (MCL2) and Rec-1 (MCL3). All three cell lines are all Cyclin D1 positive due to the hallmark t(11;14) translocation event, the initiating lesion in the majority of MCL patients (25,55). A heatmap of differentially expressed genes among all three MCL cell lines reveals that there are differences in transcript expression in all three cell MCL lines, with little variability within duplicate samples (A versus B) for each cell line (Figure 1A). Some of the transcripts with high expression code for genes that are involved in drug resistance (Figure 1A and Supplementary Table 1). Among the three cell lines, Rec-1 has the highest levels *ABCB1, ABCA6 and ABCB7*, while Jeko-1 has the highest levels of the drug efflux transporter *ABCG1.* The Granta-519 cell line express the highest levels of *ABCB9*, the second highest levels of *ABCA6* as well as the highest levels of the glycoprotein *SDC1* that has been implicated in chemoresistance (56). Gene Ontology (GO) enrichment analysis showed that for annotations of molecular function (MF) the most significant processes was protein binding (Figure 1B and Supplementary Figure 1). Other significantly enriched MF processes were those for RNA binding, DNA binding, ATP binding, metal ion binding, protein transport and transferase activity (Figure 1B and Supplementary Figure 1). Biological processes (BP) that were significant included those involved in transcription, the cell cycle, DNA damage stimulus, DNA repair and cell division (Figure 1B and Supplementary Figure 1.

**Figure 1.**
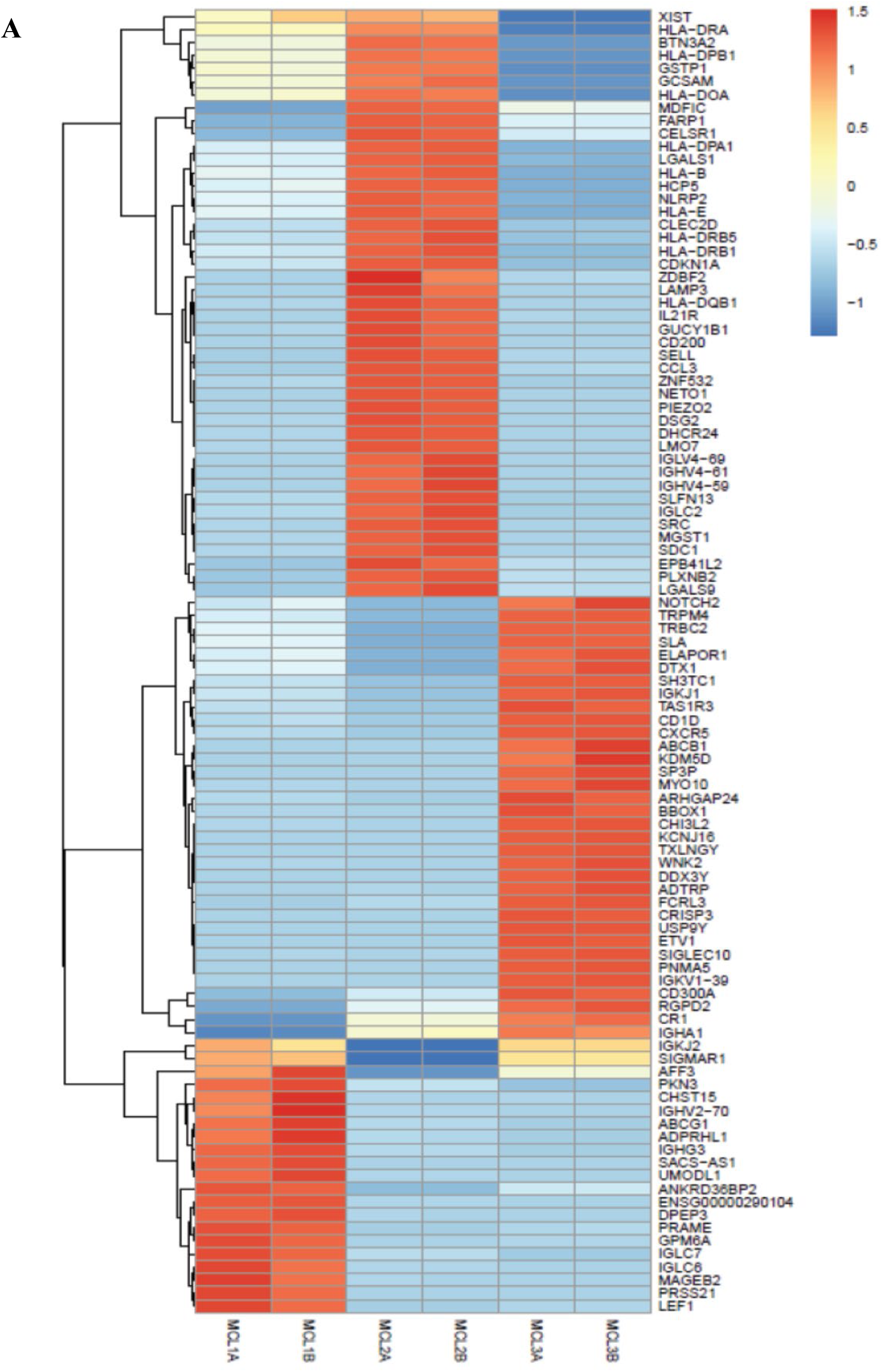
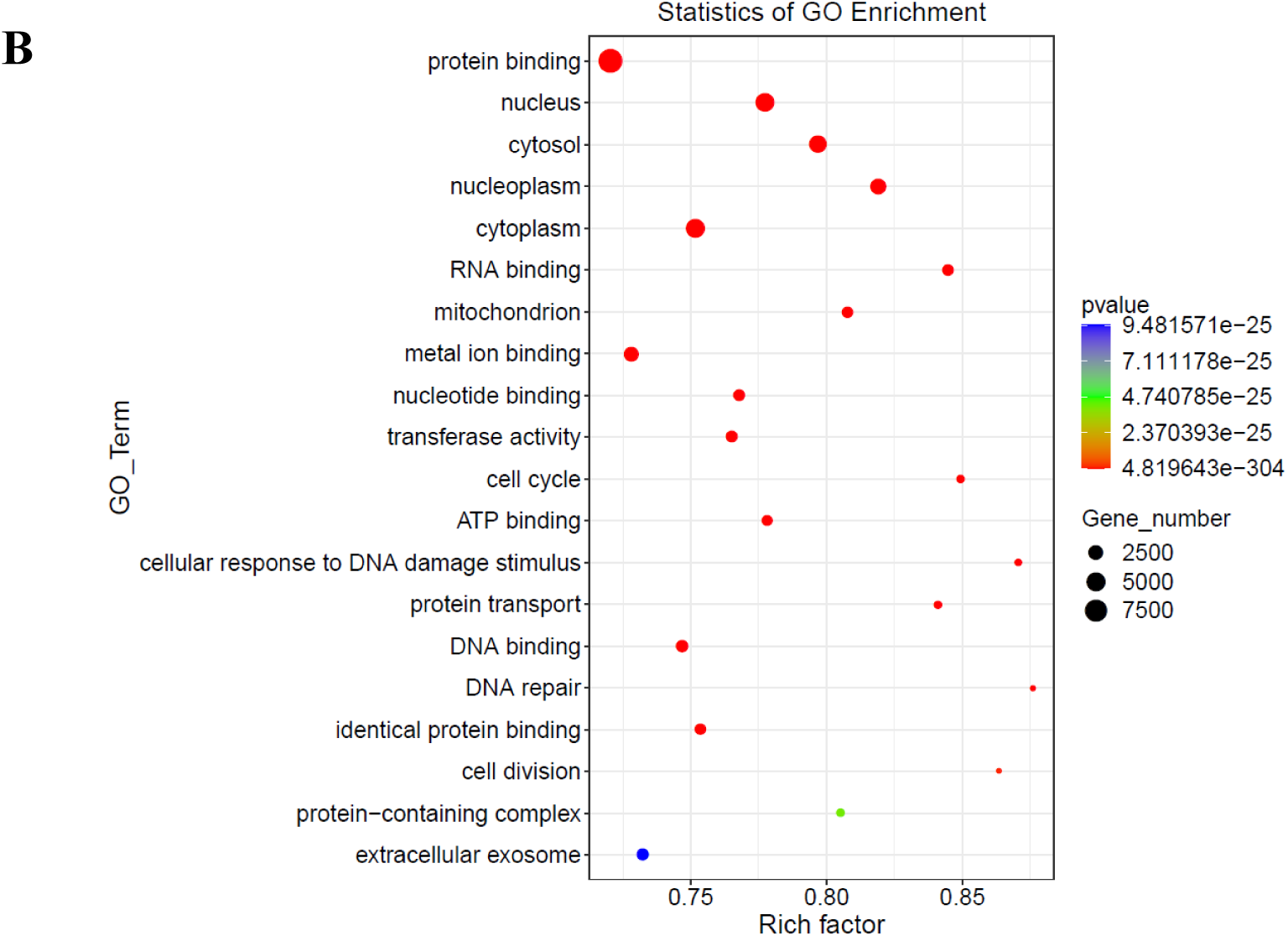
Differential transcript expression and enrichment analysis. **(A**) Heatmap showing changes in the top differentially expressed genes in three mantle cell lymphoma (MCL) cell lines in duplicate (Jeko-1: MCL1A and MCL1B, Granta-519:MCL2A and MCL1B, and Rec-1:MCL3A and MCL1B. (**B)** Gene Ontology (GO) enrichment of terms in MCL cells. Different colors show different p-values, and the size of the bubble represents the proportional gene ratio of the GO term.

### Impact of alternative polyadenylation (3’UTR-APA) on the expression of select transcripts

While the *CCND1* transcript is known to undergo APA within the same terminal exon (3’UTR-APA), not much is known about the role of 3’UTR-APA on the expression of other genes that contribute to mantle cell lymphoma oncogenesis. This includes the transcription factor, *SOX11*, another diagnostic biomarker of MCL in addition to *CCND1* (57). While *SOX11* regulates several developmental processes, it is typically absent in adult differentiated tissues including normal B-cells; however, it is highly expressed in over 90% of neoplastic MCL where it plays an oncogenic role including increasing tumor growth (57,58).

The *SOX11* 3’UTR is 7338 nucleotides long (Supplementary Figure 2A) and contains multiple potential polyadenylation signals (PASs) that can result in transcripts with 3’UTRs of different lengths. The predominant transcripts in normal tissues were reported to have the full-length 3’UTR (59,60). A detailed schematic showing potential polyadenylation signals in the *SOX11* transcript as predicted by polyA_DBv4 (https://exon.apps.wistar.org/polya_db/v4/) is provided (Supplementary Figure 2A). We developed several forward and reverse primers located in different regions of the open reading frame and the 3’UTR for *SOX11* (RefSeq transcript NM_003108) as shown (Figure 2A, Supplementary Figure 2A). To verify expression of *SOX11* in MCL cells, we performed qRT-PCR on a panel of MCL cell lines and found that all the MCL cell lines and hIPSCs expressed high levels of *SOX11* which was undetectable in both normal B-cells and the RPMI-1788. (Figure 2B). We also used primers targeted to the distal region to determine the relative use of the long 3’UTR and the short 3’UTR using qRT-PCR (Figure 2C). Consistent with a failure to detect SOX11 transcript in RPMI-1788, we also did not detect usage of the distal UTR, while there is variation in the usage of the distal UTR in the MCL cell lines.

**Figure 2.**
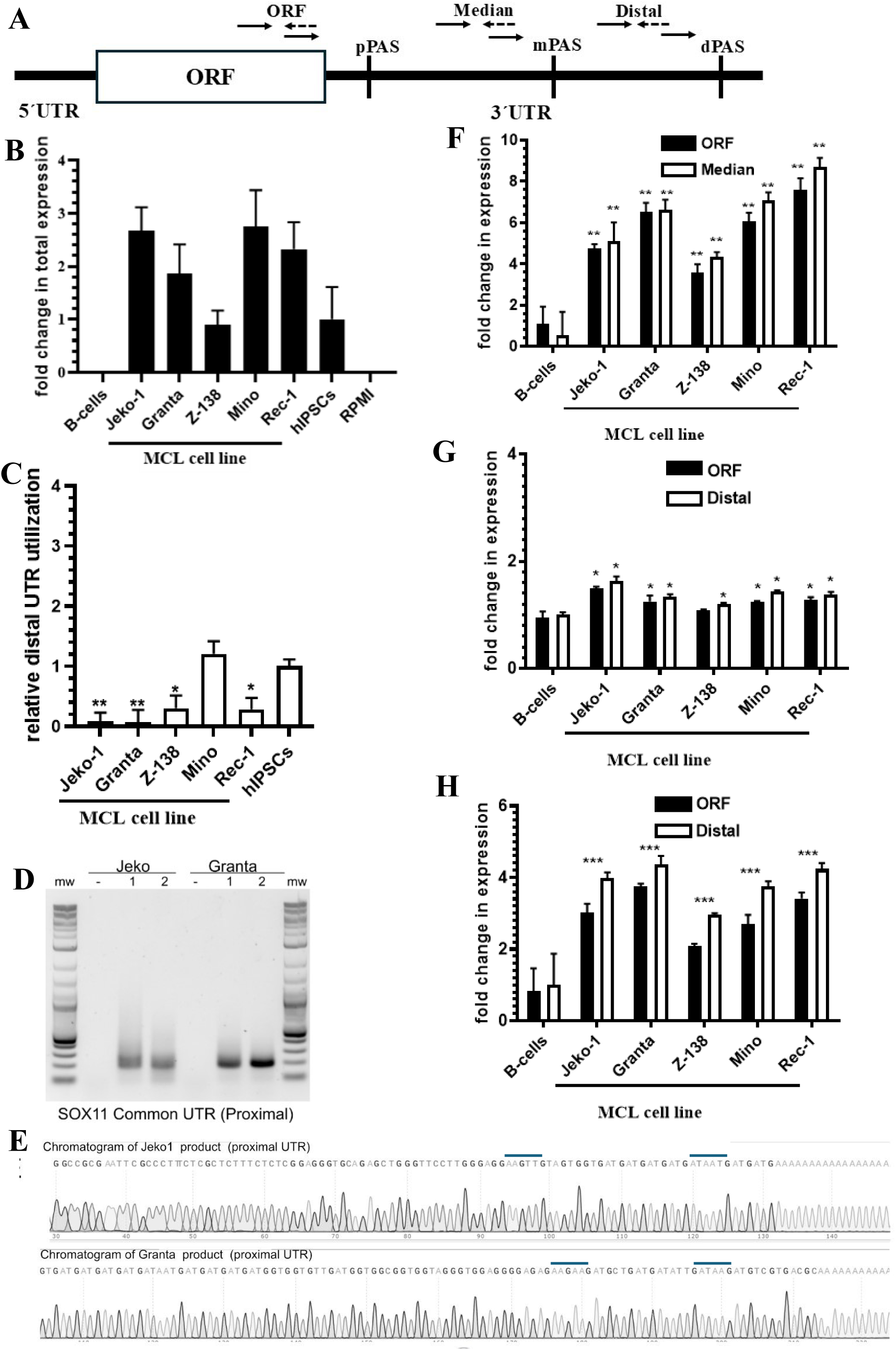
Expression of different transcripts. **(A)** Schematic representation showing the relative location of primers designed to target the open reading frame (ORF) and the 3’UTR for transcripts with approximate locations of proximal (pPAS), median (mPAS) and distal (dPAS) polyadenylation sequences. **(B)** Quantification of the SOX11 transcript in Mantle Cell Lymphoma cell lines, normal B-cells and human induced pluripotent cells, showing relative expression normalised to hIPSCs, n=3. Data is shown as the mean +/- standard deviation from the mean. **(C)** Quantification of the distal UTR usage of SOX11 in mantle cell lymphoma cell lines using modified ΔΔΔCT, normalised to hIPSCs. n=3, except for Mino where n=2. Data is shown as the mean +/- standard deviation from the mean and *P<0.01,**P<0.001. Data is from One Way ANOVA with Dunnett’s Multiple Comparisons test. **(D)** PCR products of the nested PCR of the 3′RACE resolved on 1% agarose gel in 1X TAE Buffer. Primers are targeted near the stop codon of SOX11 to amplify all the UTR transcripts. **(E)** Chromatogram of sequences of the SOX11 products cloned and amplified from the 3’RACE of Granta-519 and Jeko-1. qRT-PCR results of genes showing fold-change in expression of **(F)** *ATM*, **(G)** *BAX* and **(H)** *BTK* in different mantle cell lymphoma cell lines compared to expression in B-cells after normalizing to GAPDH control, n=3. Data is shown as the mean +/- standard deviation from the mean. *P<0.001 **P<0.0001 from two-way ANOVA with Dunnett’s multiple comparisons test compared to B-cell open reading frame (ORF), median 3’UTR or distal 3’UTR.

To identify the PAS used by *SOX11* in MCL we performed 3’rapid amplification of cDNA ends (3’RACE) on two MCL cell lines Jeko-1 and Granta-519 (Figure 2D). Next, the 3’RACE products were cloned and sequenced using Sanger Sequencing. The 3’RACE results confirmed that that the *SOX11* transcripts in Granta-519 and Jeko-1 cells primarily had short 3’UTRs (Figure 2D). We were unable to amplify *SOX11* transcripts with the longer full-length 3’UTRs (not shown). For 3’end formation, pre-mRNA processing involves cleavage downstream of a polyadenylation signal (PAS). The most prevalent hexameric PASs are AAUAAA followed by AUUAAA (61). While the Sanger Sequencing results identified the poly(A) tail for each cell line, we were unable to detect any canonical PAS upstream of the poly(A) tail (Figure 2E). To identify the potential PAS signals that were used we looked at least 10 nucleotides upstream of the poly(A) tail and searched for hexanucleotide motifs that start with the nucleotide A and contain at least two A’s. The PAS’s used to generate short 3’UTRs for SOX11 in Jeko-1 cells were AAGTTG or ATAAAT, and ones used for Granta-519 were AAGAAG or ATAAGA.

Following the same concept we also designed site specific primers for *ATM, BTK* and *BAX* (Figure 2A). All three have 3’UTRs of different sizes. The Ensembl target transcripts used were for *ATM* (ENST00000452508.7, 3’UTR length 3,594nt), *BTK* (ENST00000621635.4, 3’UTR length 438nt) and *BAX* (ENST00000293288.12, 3’UTR length 701nt). We did not detect any PCR products within the distal 3’UTR of the *ATM* transcript (not shown) so we designed transcripts within the median portion of the 3’UTR instead. We verified the presence of *ATM* in both cell lines using two different primer pairs within the open reading frame (O-1 and O-2) and the median 3’UTR region (M-1 and M-2) using PCR and running the products on an agarose gel (Supplementary Figure 2B). Using a similar approach for *BAX* (Supplementary Figure 2C) and *BTK* (Supplementary Figure 2D), we were able to validate our primers and detect transcripts using primers targeting both the open reading frame (O1 and O-2) and the distal 3’UTR region (D-1 and D-2).

To quantify potential differences in transcript levels due to 3’UTR-APA expression, we used primers targeting different regions for qRT-PCR. All the mantle cell lymphoma cell lines expressed at least a three-fold higher levels of *ATM* when compared to B-cells for each differently targeted primer set (Figure 2F). Levels of *BAX* were about 1.5 fold higher in the cancer cell lines than the B-cells (Figure 2G), while those for *BTK* were at least two-fold higher (Figure 2H). However, there did not appear to be any significant changes in 3’UTR-APA when expression was measured using the primers targeting the ORF or the median/distal targeted primers for all three genes (Figure 2F, G, and H).

In order to identify the PASs used by *ATM*, *BTK* and *BAX* within the terminal 3’UTR, we used 3’RACE. Nested forward primers within the open reading frame (O-3’R) were used to amplify transcripts that could result from 3’end cleavage and polyadenylation using a PAS proximal to the open reading frame (O-3’R). Nested forward primers were also used to amplify sequences within the median region of the 3’UTR (M-3’R) or the distal region of the 3’UTR (D-3’R). After 3’RACE, the samples were run on agarose gels. For *ATM*, we observed two differently sized products in Jeko-1 and only one product in Granta-519 using the open reading frame nested primers (O-3’R) (Figure 3A). Only one product was detected using primers within the median 3’UTR. Sanger Sequencing of the different sized *ATM* 3’UTR products gave the same sequence and identified the PAS used as AATAAA, a canonical sequence located within the median region of the 3’UTR for both cell lines (Figure 3B). Two different sized products were also observed for *BAX* in Jeko-1 using the primers in the open reading frame (O-3’R) while Granta-519 had only one (Figure 3C). One product was observed using the primers in the distal 3’UTR (D-3’R)(Figure 3C). All the products when sequenced gave the same identical sequences and the PAS, AATAAA, which was the most distal PAS within the 3’UTR (Figure 3D). The results for *BTK* were similar with the sequenced products all identifying the same canonical distal PAS within the 3’UTR (Figures 3E and 3F). To look at global PAS usage within the 3’UTR, we analyzed the RNA-Seq data of all MCL cell lines and identified the most predominantly used PASs as AATAAA (29%) and AGCGAG (29%) followed by AGCAAG (25%), and AAUAGA (3.2%) (Figure 4). Other PAS sequences made up 12.9%.

**Figure 3.**
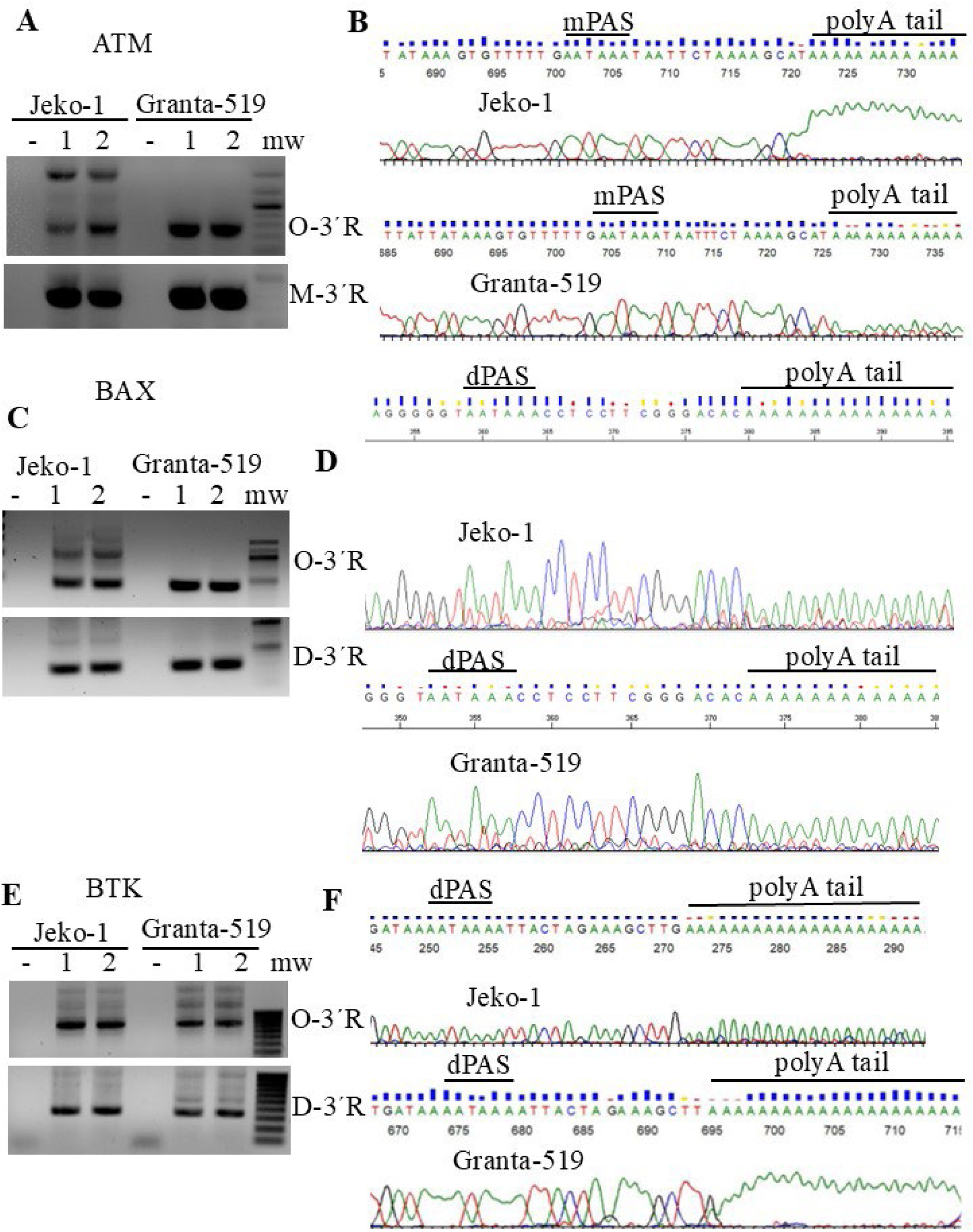
Identification of polyadenylation site usage for select transcripts. Ethidium-Bromide gels show the 3’RACE amplification products of **(A)** *ATM*, **(C)** *BAX*, and **(E)** *BTK* in mantle cell lymphoma cell lines. Products using forward nested primers within the open reading frame (O-3’R), median portion of the 3’UTR (M-3’R) and the most distal portion of the 3’UTR (M-3’R). Sanger Sequencing data showing the polyadenylation (PAS) signal used and the poly(A) tail for **(B)** ATM, **(D)** BAX and **(F)** BTK. Only the partial sequences are shown for median polyadenylation site (mPAS) distal polyadenylation site (dPAS) usage.

**Figure 4.**
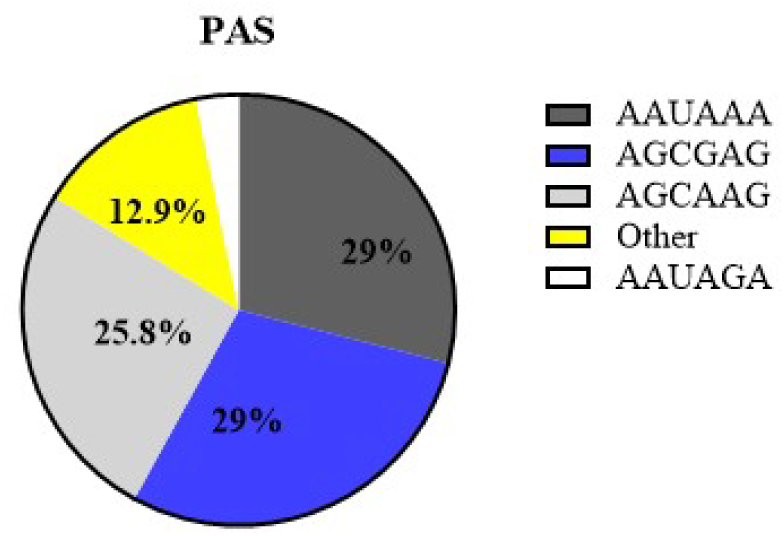
Usage of polyadenylation signals (PASs) in mantle cell lymphoma. The most frequently used PASs used are shown.

For a global comparison of 3’UTR-APA, we used DaPars2 (41,42) which measure percentage change of distal poly(A) site usage index (PDUI) on the three MCL cell lines and found no significant differences (p>0.05) in the size of the 3’UTRs (Supplementary Table 2) among the three cell lines. Generation of either short 3’UTRs (PDUI values near 0) or long 3’UTRs (PDUI values near 1) was similar in all cell lines (Supplementary Table 2). Due to the challenges of obtaining and successfully culturing normal human B-cells to obtain high quality RNA suitable for sequencing, we compared our data from MCL cells with publicly available data from normal human CD19+ B-cells. These cells were isolated using negative magnetic cell sorting to avoid changes in gene expression which may result from other cell isolation methods (62). The PDUI values were similar for most of the transcripts regardless of whether they were from the mantle cell lymphoma cell lines or the normal B-cells (Supplementary Table 3). When we analyzed the read density of the RNA-Seq data we did not see any notable differences in the 3’UTR read density profiles for *ATM*, *BAX*, *BTK and SOX11* (Supplementary Figure 3A,3B,3C and 3D). Notably, *SOX11* was only detected in the Jeko-1 and Granta-519 cell lines and was generally undetectable in normal B-cells (Supplementary Figure 3D).

### Characterization of alternative splicing of transcripts from RNA-Seq

In order to identify differences in alternative splicing we initially compared our RNA-Seq duplicate samples from the Jeko-1 (Cell line1) and the Granta-519 (Cell line2) MCL cell lines. Our analysis showed that 64.38% of the splicing events involved skipped exons(SE), 15.45% involved retained introns (RI) while 7.72% and 7.47% involved alternative 3’(A3SS) and 5’(A5SS) splice sites respectively (Figure 5A). To look at alternative exon usage in both cell lines, we used Sashimi plots to look at exons on select genes located at specific genomic coordinates provided (63). Tumor Protein Translationally-Controlled 1 (*TPT1)* is ubiquitously expressed and has multiple roles in cancer and other diseases (64). There were differences in exon usage between the two cell lines for *TPT1* (Figure 5B). Alternative splice isoforms of the arginine methyltransferase, *CARM1 (PRMT4)* (65) have different subcellular localizations and enzymatic activity in cancer (reviewed (66)). The results for *CARM1* show that while there is similarity in splicing between the two cell lines, Granta-519 cells have more transcripts than Jeko-1 cells and they preferentially use a different exon than Jeko-1(Supplementary Figure 4A). Both cell lines expressed low levels of carnitine palmitoyl transferase 1B (*CPT1B),* and alternative exon usage was also observed for this gene (Supplementary Figure 4B). The expression of *CPT1B*, which is involved in fatty acid metabolism whose variable expression, has been linked with various types of solid cancers as well as leukemia (reviewed (67)).

**Figure 5.**
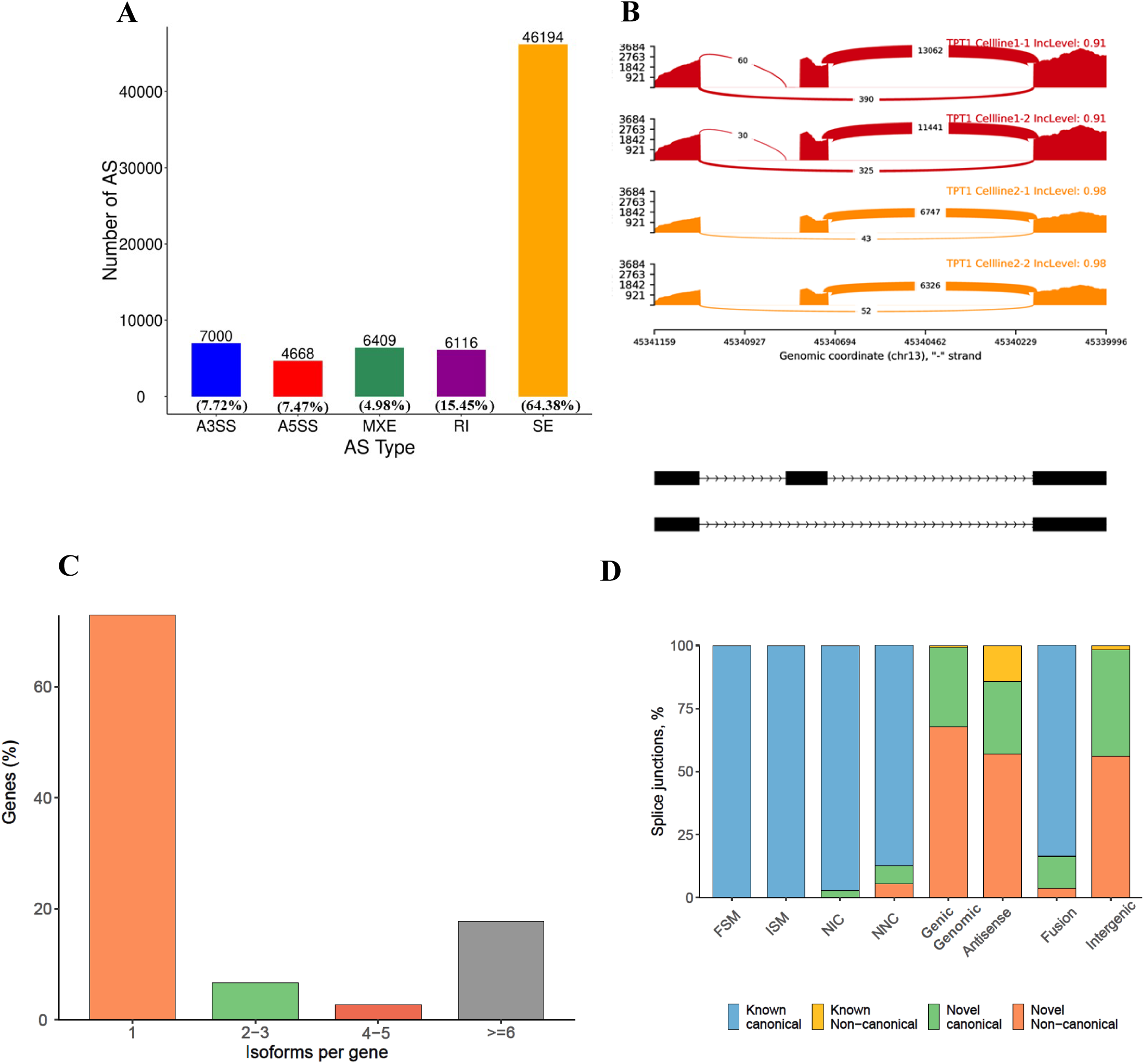
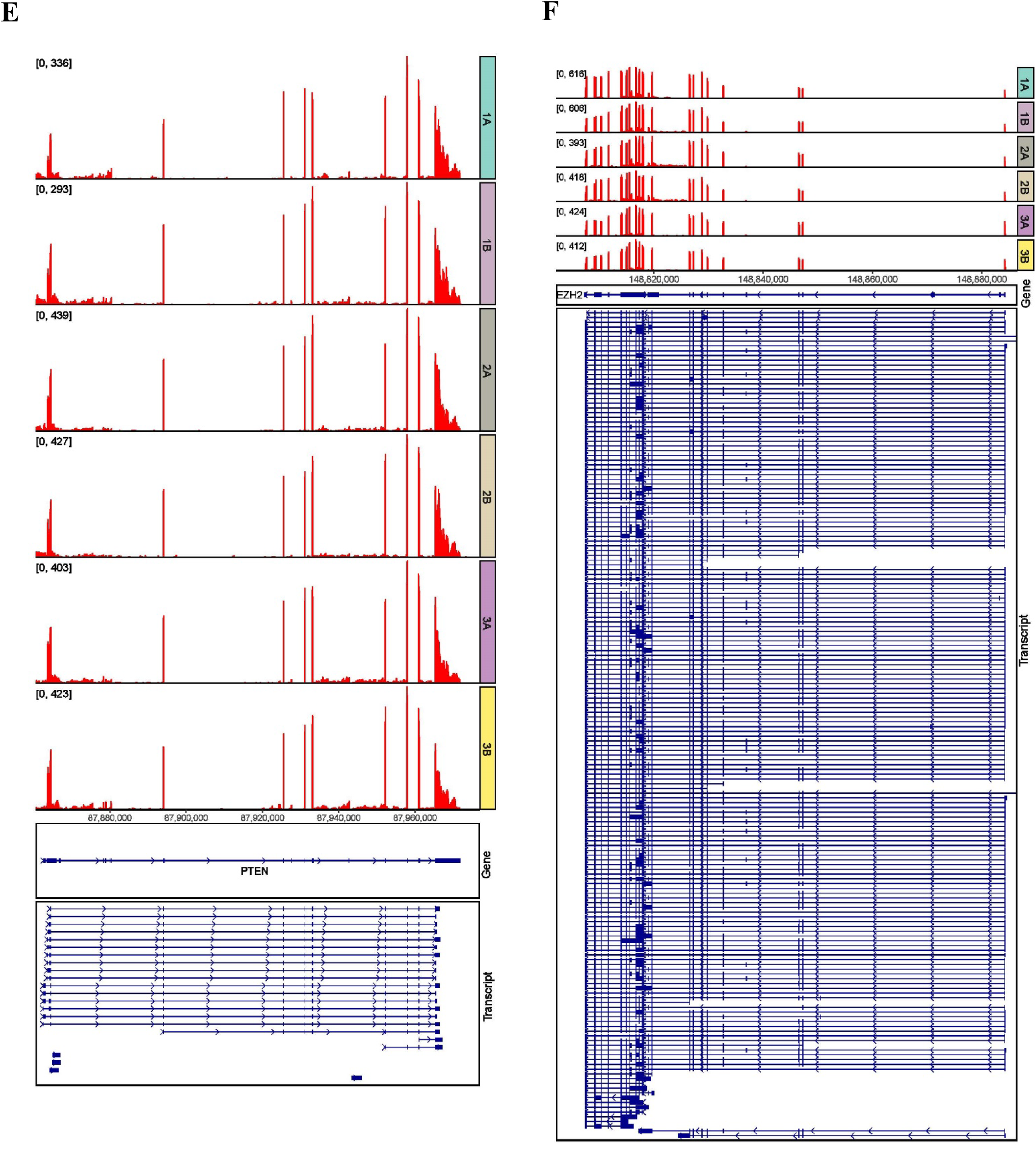
Alternative splicing in mantle cell lymphoma. **(A)** Short-read sequencing data distribution of five different types of alternative splicing. (A3SS-Alternative 3’Splice Site, A5SS-Alternative 5’Splice Site, MXE-Mutually Exclusive Exons, RI-Retained Intron, SE-Skipped Exon). **(B)** Sashimi plot of short-read RNA Sequencing data showing model IGV views of exon skipping (SE) events for *TPT1*. Comparisons are between duplicate samples of two mantle cell lymphoma cell lines Jeko-1 (Cell line 1-1 and 1-2) and Granta-519 (Cell line 2-1 and 2-2) with annotated exons shown underneath. **(C)** SQANTI3 identification of isoforms from long-read Iso Sequencing data. **(D)** SQANTI/SQANTI3 splice junction based transcript categories: full-splice match (FSM); incomplete-splice match (ISM); novel-in-catalog (NIC); novel-not-in-catalog (NNC);genic genomic; antisense; fusion; and intergenic. (50,68). Isoform diversity showing different transcripts for **(E)** *PTEN* and **(F)** *EZH2* as supported by RNA Sequencing reads (top panels) and Iso Sequencing reads in all three MCL cell lines in duplicate. Shown is Jeko-1 (1A and 1B), Granta-519 (2A and 2B) and Rec-1 (3A and 3B).

### Characterization of alternative splicing of transcripts using Iso-Seq and RNA-Seq

We also had long-read RNA Iso Sequencing (Iso-Seq) data from all the three MCL cell lines (Jeko-1, Granta-519 and Rec-1) generated by our lab (48), Accession number E-MTAB-17128 (ArrayExpress (BioStudies) database (http://www.ebi.ac.uk/arrayexpress). Our analysis of the Iso-Seq data found that the majority of genes had one predominant isoform and surprisingly, almost 15% of genes had six or more isoforms (Figure 5C) (50,68). When we looked at the splice junctions based on the different SQANTI3 structural categories we found that most splice junctions are canonical followed by novel non-canonical splice junctions (Figure 5D) (50,68). For a more comprehensive look at transcript diversity in MCL we combined our RNA-Seq data with the Iso-Seq data for all three cell lines. There were multiple transcripts for the tumor suppressor *PTEN* (Figure 5E). There was a surprisingly large number of alternative transcripts for *EZH2* (Figure 5F) whose clinical expression is associated with reduced survival of MCL patients (69). We found 11 different transcripts for *NDRG1* which has 17 exons and has been associated with different types of cancer (70) (Supplementary Figure 5A). There were 13 transcripts for *MAP2K7* including the isoform *MAP2K7* Δexon 2 which is associated with dedifferentiation (Supplementary Figure 5B)(71).

### Alternative splicing isoforms of glutaminase (GLS1) and their susceptibility to GLS1 inhibitors

For functional validation of different transcripts, we investigated the expression and impact of targeting glutaminase (*GLS/GLS1*) in MCL. Since MCL is incurable due to drug resistance and a high recurrence rate, there is a lot of interest therapeutically targeting drug resistant disease. While MCL patients initially respond to Ibrutinib, a Bruton tyrosine kinase (*BTK*) inhibitor, patients often develop resistance to the drug, and this has been associated with increased *GLS1* expression and increased glutamine metabolism in cancer (72). *GLS1* is the first enzyme involved in glutaminolysis and has one main splice variant GAC that is important for cancer metabolism (73). The Iso-Seq data showed that there are several splice variants for *GLS1* in all three MCL cell lines (Figure 6A). We were also able to detect the two major splice isoforms, GAC (exons 1-15) and KGA (exons 1-18 with an alternative exon 15) in the Iso-Seq data and this was supported by the RNA-Seq data (Figure 6B). We used qRT-PCR to quantify GLS isoforms using primers located in the regions shown that were specific for the KGA and GAC 3’UTRs as well as within the open reading frame of both isoforms (Figure 6B). All the MCL cell lines expressed higher levels of the KGA transcript than the normal B-cells with the highest expression in the Rec-1 cell line(Figure 6C). The Jeko-1 and Rec-1 cell lines expressed high levels of the GAC transcript (Figure 6D). To determine expression at the protein level we used Western blotting (Figure 6E). When normalized to the protein levels in B-cells, the Mino cell line expressed the highest levels of GAC protein followed by Z-138 and Jeko-1. The levels of GAC protein in Granta-519 and Rec-1 were similar and slightly higher than B-cells. The Mino cell line is the only one that expressed higher levels of the KGA protein than B-cells.

**Figure 6.**
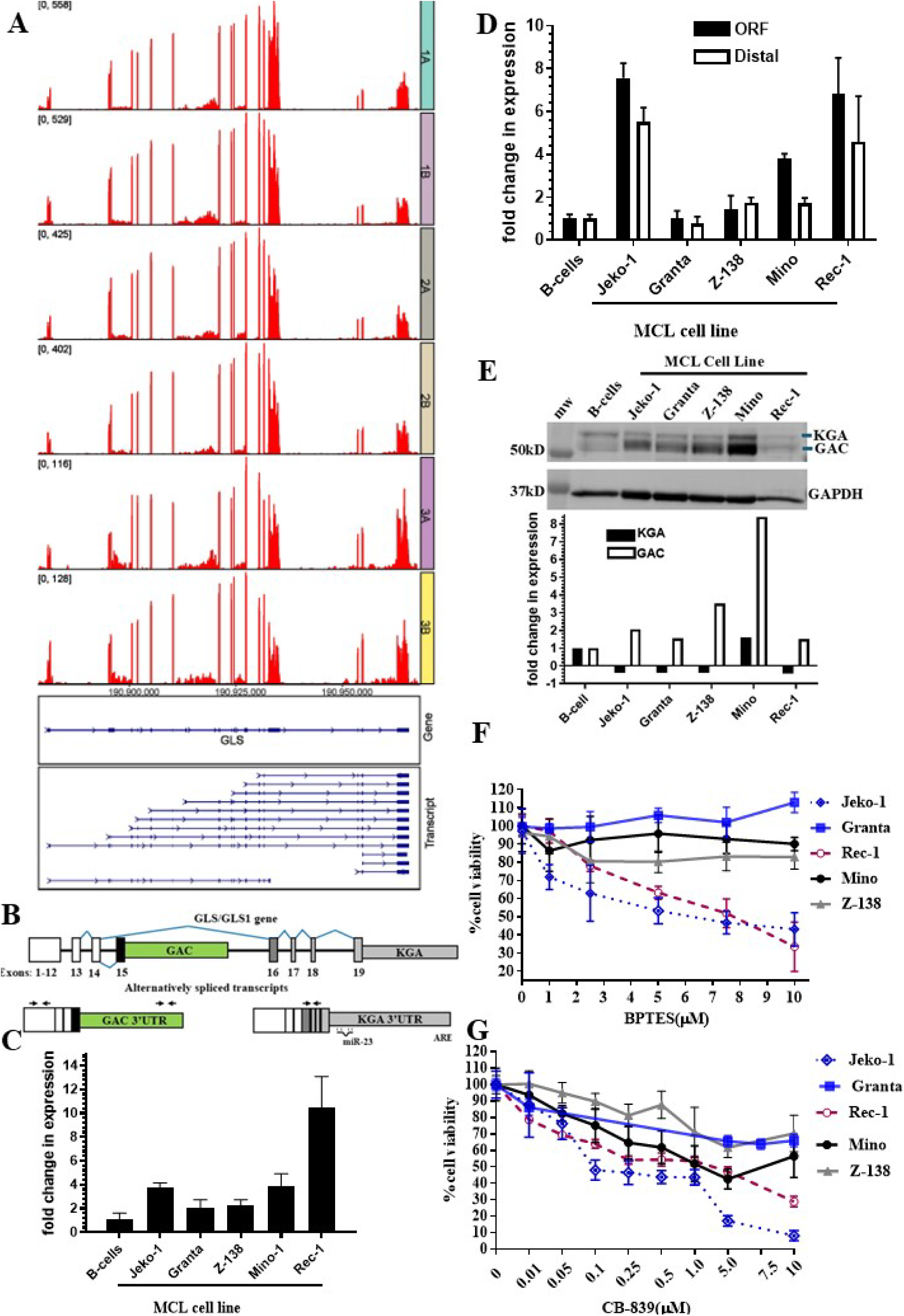
Alternative splicing of glutaminase (GLS1) and effects of GLS1 inhibitors on cell viability. **(A)** An overlay of RNA-Sequencing and Iso Sequencing reads for GLS1 in three different mantle cell lymphoma cell lines in duplicate Jeko-1 (1A and 1B), Granta-519 (2A and 2B) and Rec-1 (3A and 3B). **(B)** A schematic representation of the GLS1 gene (top panel) and the two alternatively spliced isoforms (GAC and KGA). The approximate locations of the miR-23 target site and the AU-rich (ARE) element are also shown within the KGA isoforms 3’UTR. **(C)** qRT-PCR results showing fold change in levels of the KGA isoform in different cell lines compared to levels in B-cells after normalizing to GAPDH. **(D)** qRT-PCR results showing fold change in levels of the GAC isoform using two sets of primers one set located close to the open reading frame (ORF) and the other towards the distal end of the 3’UTR (Distal) (E) Western blot analysis of cell lysates from different cell lines probed using an antibody that detects both the GAC and KGA isoforms (top panel). Quantification of the Western blot was done for the GAC and KGA isoforms and results were plotted as fold change when compared to levels in B-cells (bottom panel). Cells were treated with an increasing concentration of the GLS1 inhibitors BPTES **(F)** and CB-839 **(G)** for 48 hours. Cell viability for each concentration was compared to cells treated with the delivery vehicle DMSO using MTT assays. The cell viability in DMSO (0 µM drug concentration) was set to 100% (n=4).

In order to determine the effect of glutaminase inhibition on cancer cell viability, we treated cells with increasing concentrations of two glutaminase inhibitors, the classical prototype bis-2-(5-phenylacetamido-1,2,4-thiadiazol-2-yl)ethyl sulfide (BPTES) and its more potent analog, CB-839 (74) which target both KGA and GAC. Our results show that that the Rec-1 and Jeko-1 cell lines had the greatest decrease in cell viability when treated with BPTES(Figure 6F). Z-138 cells showed intermediate decrease in cell viability, while Granta-519 and Mino did not see significant effects from BPTES treatment. All the cell lines were affected by treatment with CB-839 at the highest concentration used (Figure 6G). Similar to BPTES, Rec-1 and Jeko-1 cells showed the greatest decrease in cell viability after treatment with CB-839 (Figure 6G).

## DISCUSSION

Using both short-read RNA-Seq and long-read Iso-Seq data, in this study we analyzed global patterns of alternative polyadenylation and alternative splicing using the blood cancer mantle cell lymphoma (MCL), that originates from B-cells, as our experimental model. Due to its biological and molecular heterogeneity, MCL is no longer considered a single disease, and its classification and subtyping are constantly being updated based on emerging data (33,54). A major focus of research in MCL has focused on identifying mutations in driver genes, and there is not a lot of information on alternative splicing and alternative polyadenylation in MCL. The findings in this study provides a potential blueprint on using a combination of different sequencing data to characterize alternative transcripts generated in different transcriptomes.

### There is heterogeneity in gene expression

In order to determine similarities and differences, we initially used three different MCL cell lines (25,55). Interestingly, when we compared gene expression in all three cell lines, the heatmap showed differences as well as some overlaps in the top expressed genes in each cell line. Hence, heterogeneity can arise within the same classical MCL subtype (33,54). More work still remains to be done on the biological/clinical relevance of these variations.

A notable observation from our work showed that each of the three cell lines had higher levels of mRNA expression of genes associated with drug resistance compared to the other two cell lines. Two of the genes, *ABCB1* which codes for P-glycoprotein, also known as the multidrug resistance protein 1, and *ABCG1,* are well-known ABC drug efflux transporters. Apart from chemoresistance they have both been linked to other hallmarks of cancer including increased cancer stemness (as reviewed (75)). One cell line had upregulated levels of *SDC1* which is associated with chemotherapy drug resistance (56). These differences in MCL cell line data suggest that different genes may contribute more to drug resistance than others in different MCL patients.

Consistent with what is known, our Gene Ontology (GO) analysis picked up several significant pathways that have long been associated with MCL pathogenesis including the cell cycle, and DNA damage response (76).

### Formation of the 3’ends and alternative polyadenylation is not solely dependent on the canonical polyadenylation signal for most genes

The length of the 3’UTR affects the sequences that are located within the 3’UTR including miRNA binding and RNA binding protein regions. This subsequently affects the stability, translational efficiency, and function of the entire mRNA transcript and is associated with disease (21). Elements that determine the size of the 3’UTR include the polyadenylation signal (PAS), and in mammals the canonical sequence AAUAAA is frequently found upstream of the 3’end cleavage and polyadenylation site (8). Besides *CCND1* (30), not much is known about the PASs used for genes associated with MCL, and global 3’UTR-APA patterns in MCL are still unknown.

In addition to *CCND1*, another major driver of MCL is the transcription factor, *SOX11* which is overexpressed in the majority of MCL patients (57,58). Several mechanisms contribute to the overexpression of *SOX11* in MCL including DNA methylation and histone acetylation (77,78). At the transcript level, our qRT-PCR data shows that consistent with previous reports, *SOX11* was not expressed in normal B-cells but was expressed in all MCL cell lines. While *SOX11* is constitutively overexpressed in MCL cell lines, the reported protein abundance varies (79). One way in which protein abundance can be regulated post-transcriptionally is through alternative polyadenylation (80). Our results show that in MCL, *SOX11* uses several non-optimal polyadenylations signals (PASs) resulting in generation of short 3’UTRs and shows that 3’end cleavage and polyadenylation occurring early within the 3’UTR, contrary to the predicted canonical PASs. This suggests that *SOX11* undergoes APA in MCL to eliminate generation of the longest 3’UTR containing transcripts. *SOX11* was recently identified as a novel 3’UTR-APA regulated transcript whose 3’UTR length changed during neural differentiation (81). While it was shown that the RNA-binding motif protein 3 (*RBM3*) binds in the 3’UTR of *SOX11* and regulates the mRNA stability in neuroblasts during hypothermia (82), the impact of APA on *SOX11* mRNA stability still needs further study.

When we looked at other MCL associated genes, our qRT-PCR results show that MCL cells express higher levels of ataxia telangiectasia (*ATM*), bruton tyrosine kinase (*BTK) and BAX* than normal B-cells. While overexpression of *BAX* in MCL cells has been previously reported (55), the focus for *ATM* and *BTK* has been on the frequency and impact of mutations. An estimated 30-50% of MCL patients have mutations in *ATM* (*34*). While less common, aberrant somatic mutations in *BTK* are known to be involved in MCL pathogenesis and *BTK* inhibitor drug resistance (83). The impact of these mutations on the expression levels of both genes is unknown. Next, we used 3’RACE to identify the polyadenylation signal (PAS) used in 3’end formation for each gene. While premature cleavage and polyadenylation occurring within the introns (intronic polyadenylation) has been reported for *ATM* (84), our 3’RACE results did not detect this. We found that MCL cells preferentially used a median PAS located upstream of the putative more distal canonical PAS for 3’end formation within the same 3’UTR. The *ATM* median PAS used is also a canonical sequence, AAUAAA. While 3’UTR-APA is expected for *ATM* which has a long 3’UTR (∼3,594 nt), our results detected usage of the distal PAS within the 3’UTR for both *BTK* and *BAX* which have shorter 3’UTRs, 438 and 701 respectively. The PAS used was also AAUAAA for both genes and this suggests that having an optimal PAS is important for proper 3’end processing of these transcripts in MCL cells. This is supported by previous work looking at cyclin D1 (*CCND1*) in MCL where several patient samples had mutations, deletions or duplications that generated a premature canonical PAS resulting in generation of truncated *CCND1* 3’UTRs (30). While shortened 3’UTRs resulting from 3’UTR-APA is important in evasion of miRNAs and increased mRNA stability for *CCND1* resulting in increased protein levels (85), this is not always the case for all transcripts. Hence the impact of 3’UTR-APA driven shortening of the 3’UTR for *SOX11* and *ATM* in MCL still remains undetermined. This is important because it may affect their therapeutic targeting in cancer. As an example, the cell surface glycoprotein, CD19 undergoes 3’UTR-APA which affects its expression in B-cell progenitor acute lymphoblastic leukemia (B-ALL), (86). CD19 is important because it is a molecular target for current CD19-targeted chimeric antigen receptor (CAR) T cell therapies, bispecific antibodies including Blinatumomab, and other targeted cancer therapies in different stages of development (86).

A recent study using single-cell RNA sequencing data suggests that 3’UTR-APA, within terminal exons, most frequently provides tissue-specific PAS usage (60). At a global level, when we looked at PAS usage within the terminal exons through 3’UTR-APA, we found that usage of the canonical PAS was only 29% in MCL. Contrary to expectations that the majority of the non-canonical PASs used would contain only one nucleotide permutation of the AAUAAA sequence, we found that the PAS sequences AGCGAG (29%) and AGCAAG (25.8%) were used quite frequently. Interestingly, these hexamers follow the pattern of the RRYRRR motif (R-pyrimidine C,T/U); Y-purine(A,G) that was found to occur frequently in *C. elegans* where there was no canonical PAS (87). The canonical PAS is recognized and bound by the 3’end processing factors WDR33 and CPSF30 with CPSF160 acting as a scaffold to facilitate 3’end cleavage (88). The mechanism required for 3’end cleavage of transcripts with non-canonical PASs still remains undetermined and may have other 3’end factors playing a more major role than when the canonical PAS is involved.

### Alternative splicing gives rise to numerous isoforms some with biological implications

Besides 3’end formation, alternative splicing also gives rise to different transcripts. Alternative splicing is important for the protein heterogeneity and different biological functions. The short-read RNA sequencing (RNA-Seq) data identified skipped exons (SE) as the most common in MCL which is expected since this is the most prevalent type of splicing in humans (3). Analysis of splicing patterns using long-read Iso Sequencing (Iso-Seq) data showed that in MCL, the majority of genes had one dominant splice isoform. This supports previous findings in human tissues (89). There is no consensus on the number of transcripts reported per gene (3) and determining the number of alternative transcripts per each cell type is important to fully annotate the human transcriptome.

To get a more comprehensive view of variations in splicing, we integrated the RNA-Seq and Iso-Seq data for select genes that had at least 10 transcripts. Our data identified multiple splice variants for the tumor suppressor gene, phosphatase and tensin homolog *(PTEN)*, N-myc downstream regulated gene 1 (*NDRG1*) and Mitogen-Activated Protein Kinase 7 (*MAP2K7*) and the Enhancer of Zeste Homolog 2 (*EZH2*). The histone-lysine N-methyltransferase enzyme *EZH2* is considered a master regulator of many biological processes and cancer (90). The *EZH2* gene is made up of 20 exons and different splice variants of *EZH2* have different biological effects and add an added layer regulation that affects protein expression and the gene’s role in different cancer types (91,92).

We took a more detailed look at splice variants of the metabolic enzyme, glutaminase 1 (*GLS1/GLS*). Apart from increased glycolysis, as part of their metabolic reprogramming cancer cells also increase their uptake of glutamine. The first step of glutaminolysis involves the conversion of glutamine to glutamate a rate limiting step that is catalyzed by *GLS1* (93). Our sequencing results identified several full length *GLS1* transcripts including the two main alternatively spliced isoforms, GAC and KGA. Several studies have shown that the GAC splice variant is the preferred isoform in cancer (93,94). Quantification of *GLS1* using qRT-PCR showed that while all the MCL cell lines showed higher levels of expression of the KGA transcript at the RNA level than B-cells, this did not correlate to higher KGA protein levels. With the exception of Mino cells, all the MCL cell lines had lower protein levels of KGA than B-cells. The human KGA transcript contains miR-23a/b target sites (95), while studies have shown that the rat derived KGA transcript contains a functional AU-rich destabilizing element (96). While both these are expected to result in decreased KGA stability, our qRT-PCR results showing notably high KGA transcript levels would suggest another level of KGA regulation is occurring in MCL and this may include a reduction in translation or an increase in KGA protein degradation.

Besides GLS1, cells can also express the liver-type glutaminase (LGA)/GLS2. Since GLS1 is associated with cancer, we investigated the impact of *GLS1* specific inhibitors on MCL cell lines. We found that the Granta-519 cell line which had low protein levels of both the GAC and KGA isoforms showed the least sensitivity to BPTES while the Rec-1 cell line with similar expression levels showed more sensitivity to BPTES. Mino cells, which expressed the highest levels of both GAC and KGA isoforms at the protein level, were less sensitive to BPTES. The drug resistance in Granta-519 cells may partially be due to the ability of these cells to spontaneously form 3D-speroids in cell culture which may limit cellular access to BPTES. All the cell lines show sensitivity to CB-839, a more potent analog of BPTES albeit at much higher concentrations than reported in other studies (74,97). While our research did not measure actual glutaminase enzyme activity, our results suggest that mRNA overexpression of GAC alone may need to be combined to GAC protein levels and actual GAC enzyme activity to provide a better indicator of an individual’s glutamine metabolic status and their potential response to GLS1 inhibitors. This is important because GAC has also been linked to advanced cancer and therapeutic drug resistance (74,97), and our studies have identified other genes that facilitate drug resistance in MCL.

In conclusion, we show that combining multiple sequencing techniques provides a powerful tool to investigate transcriptome diversity in normal biology and disease which will provide insights to guide further rational functional validation.

## AUTHOR CONTRIBUTIONS

Chioniso P. Masamha: Conceptualization, methodology, investigation, validation, supervision, formal analysis, visualization, writing original draft, review and editing, data curation. Mahesh Gupta: Investigation, validation, supervision, formal analysis, visualization reviewing and editing. Jie Li: Methodology, formal analysis, software and bioinformatics, visualization, writing original draft. Lael Pasipamire: Investigation, validation, formal analysis, visualization, reviewing and editing. Natoya Peart: Methodology, investigation, validation, supervision, formal analysis, visualization, writing original draft, reviewing and editing. Demiah Lockett: Investigation, validation, visualization, reviewing and editing. Jie Li: Methodology, formal analysis, software, visualization, writing original draft. Isabella Todd: Investigation, validation, formal analysis, reviewing and editing. Haavia Rajakumar: Investigation, validation, formal analysis, reviewing and editing.

## SUPPLEMENTARY DATA

Supplementary Data are available.

## DATA AVAILABILITY

The short-read (PolyA) RNA-Seq data is deposited in the Gene Expression Omnibus (GEO) and will be made available upon publication. The long-read Iso-Seq datasets are deposited in the ArrayExpress (BioStudies) database under the accession number E-MTAB-17128 (https://www.ebi.ac.uk/biostudies/arrayexpress/studies/E-MTAB-17128) (48). We also used publicly available RNA-Seq data of normal CD19+ B-cells from healthy individuals separated by negative magnetic cell sorting acquired from GEO under accession number GSE279633 (62).

