## Supplementary Figures for "Leveraging short and long-read sequencing to comprehensively characterize alternatively polyadenylated and alternatively spliced transcripts"

### Supplementary Figure Legends

**Supplementary Figure 1**. GO terms for Mantle Cell Lymphoma gene expression. The number of genes per GO-term are shown for each category .

**Supplementary Figure 2**. **Strategies to target different regions of transcripts**. **A.** Schematic of the refseq SOX11 transcript, NM_003108, showing the untranslated region (UTR) and open reading frame (ORF). Indicated by open triangles are the canonical polyadenylation signals (AAUAAA) predicted by PolyA_DB (https://exon.apps.wistar.org/polya_db/v4/) with the distance from the stop codon shown. Locations for the primers for 3'RACE indicated by arrows (not drawn to scale). Ethidium-bromide stained gels of PCR products using different primer pairs developed and validated for **B**. ATM, **C**. BAX, and **D**. BTK. Shown are PCR products obtained using primers targeting the open reading frame (O-1 and O-2) median 3'UTR (M-1 and M-2) and distal 3'UTR (D-1 and D-2).

**Supplementary Figure 3**. Comparison of RNA-Seq read density data between mantle cell lymphoma cell lines and normal B-cells. RNA-Seq read density for 3' UTR terminal exon and select exon(s) in mantle cell lymphoma cell lines, Jeko-1 and Granta-519 compared to publicly available RNA-Seq data from normal B-cells from four patients (P1-P4, Reference Henning et al 2025). Shown are the read densities for **A**. ATM, **B**. BAX, **C**.BTK and **D**. SOX11.

**Supplementary Figure 4**. **Alternative exon usage**. Sashimi plots showing alternative exon usage for select exons at specific genomic coordinates for **A**. CARM1 and **B**. CPT1B. Comparisons are of RNA-Seq data between duplicate samples of two mantle cell lymphoma cell lines Jeko-1 (Cell line 1-1 and 1-2) and Granta-519 (Cell line 2-1 and 2-2) with annotated exons shown underneath.

**Supplementary Figure 5**. **Alternative splicing.** Isoform diversity showing different transcripts for **A**. NDRG1 and **B**. MAP2K7 supported by RNA-Sequencing reads (top panels) and Iso Sequencing reads in all three MCL cell lines in duplicate. Shown is Jeko-1 (1A and 1B), Granta-519 (2A and 2B) and Rec-1 (3A and 3B) .

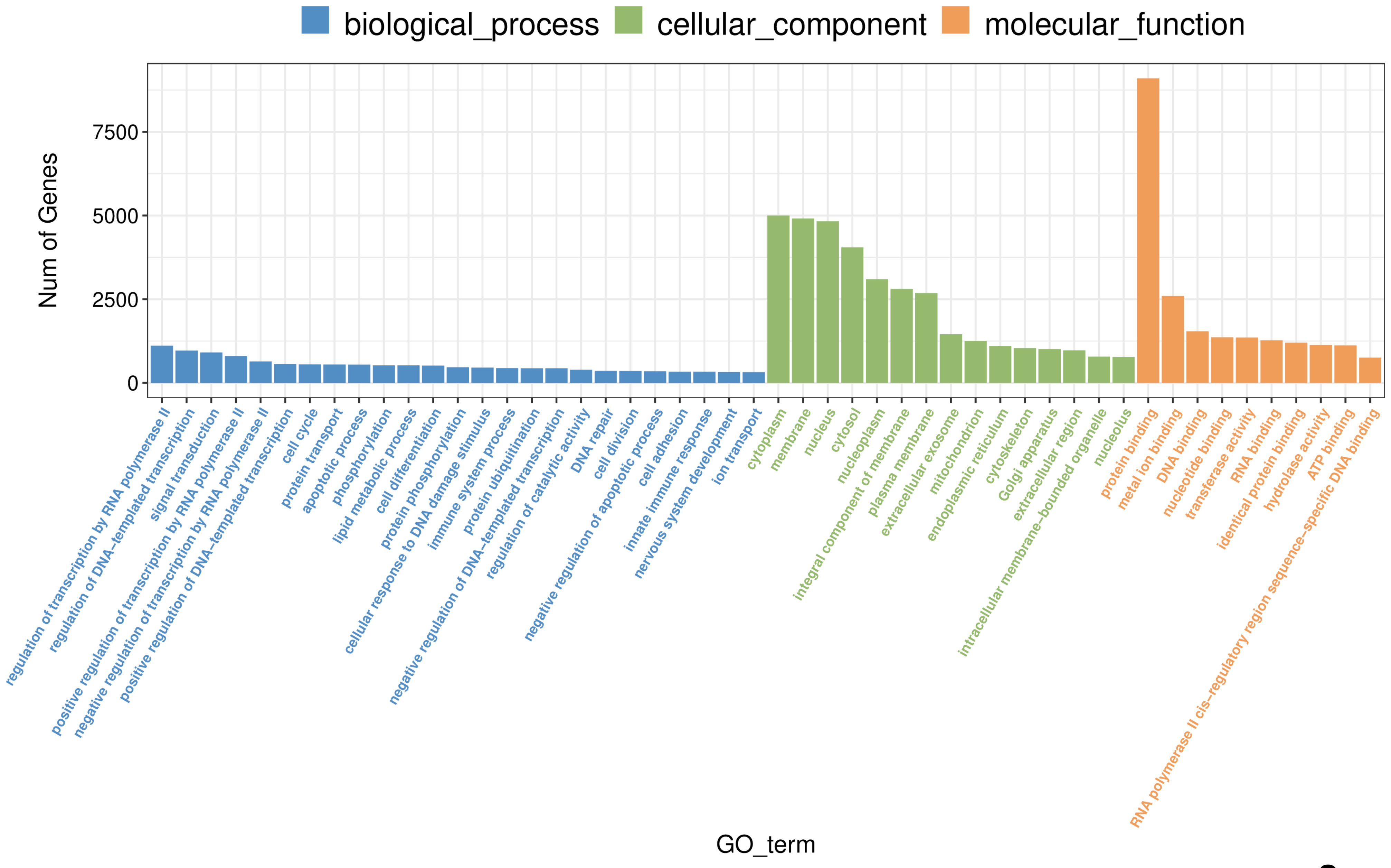


**Supplementary Figure 1**

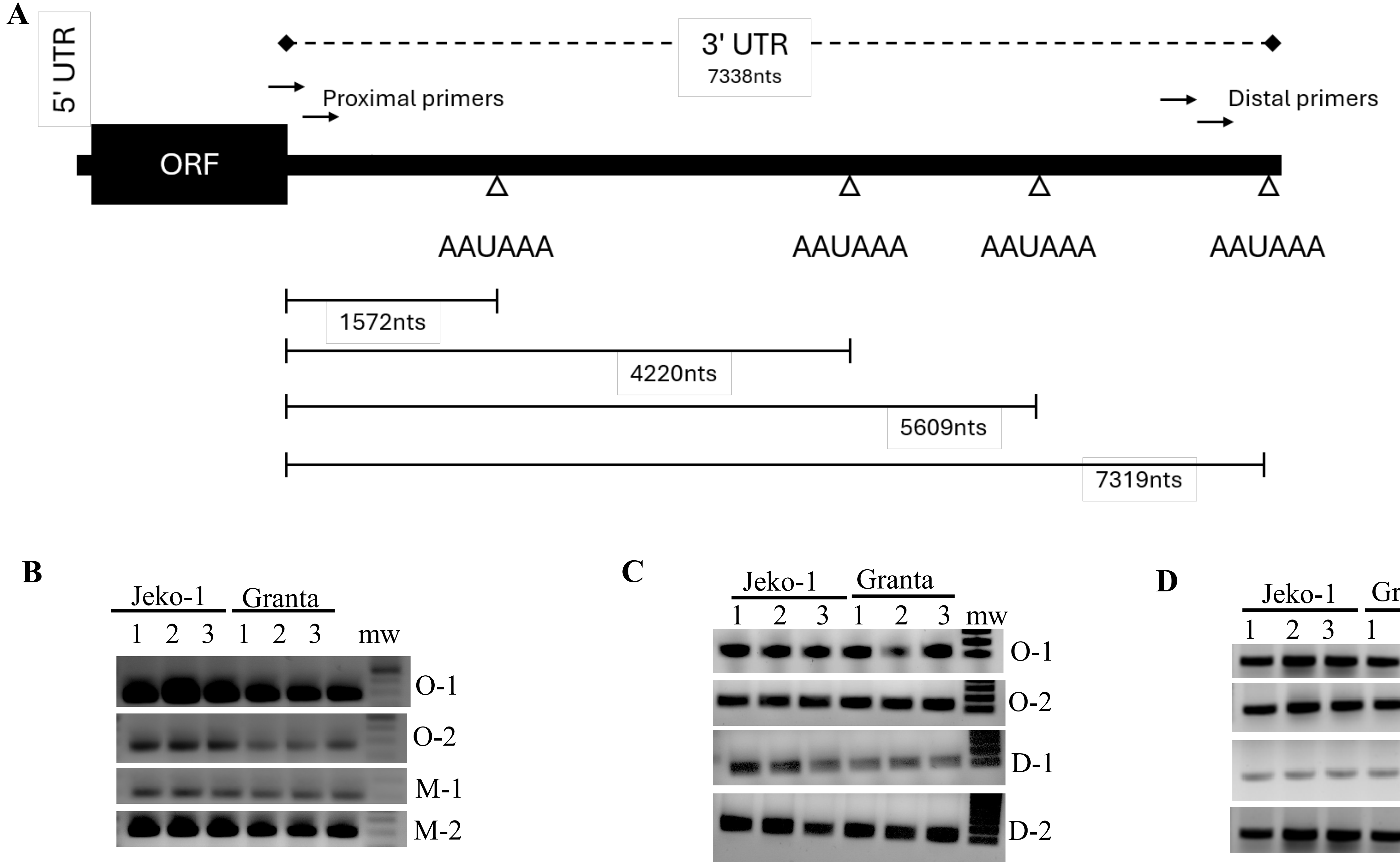


**Supplementary Figure 2**

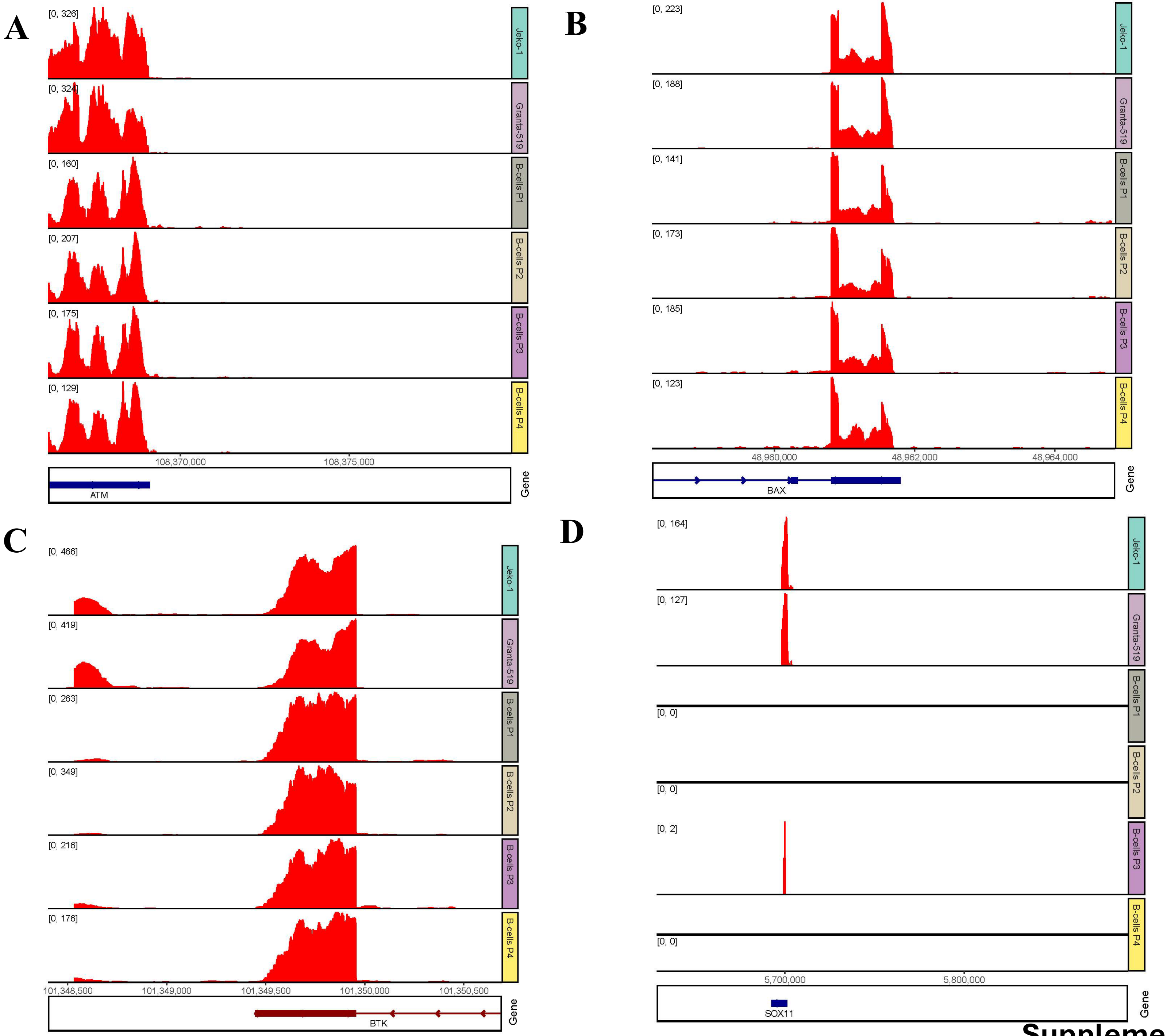


**Supplementary Figure 3**

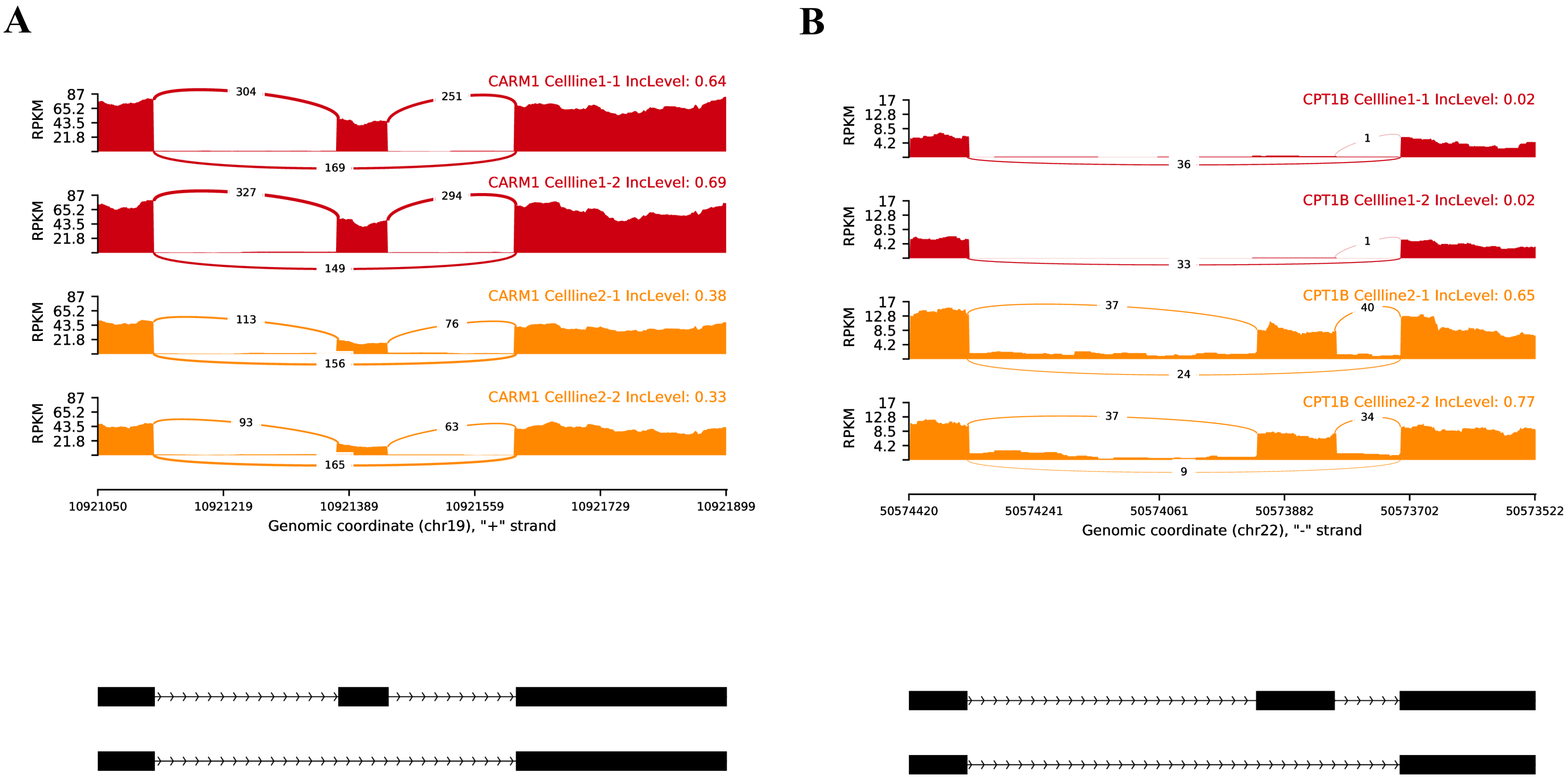


**Supplementary Figure 4**

A

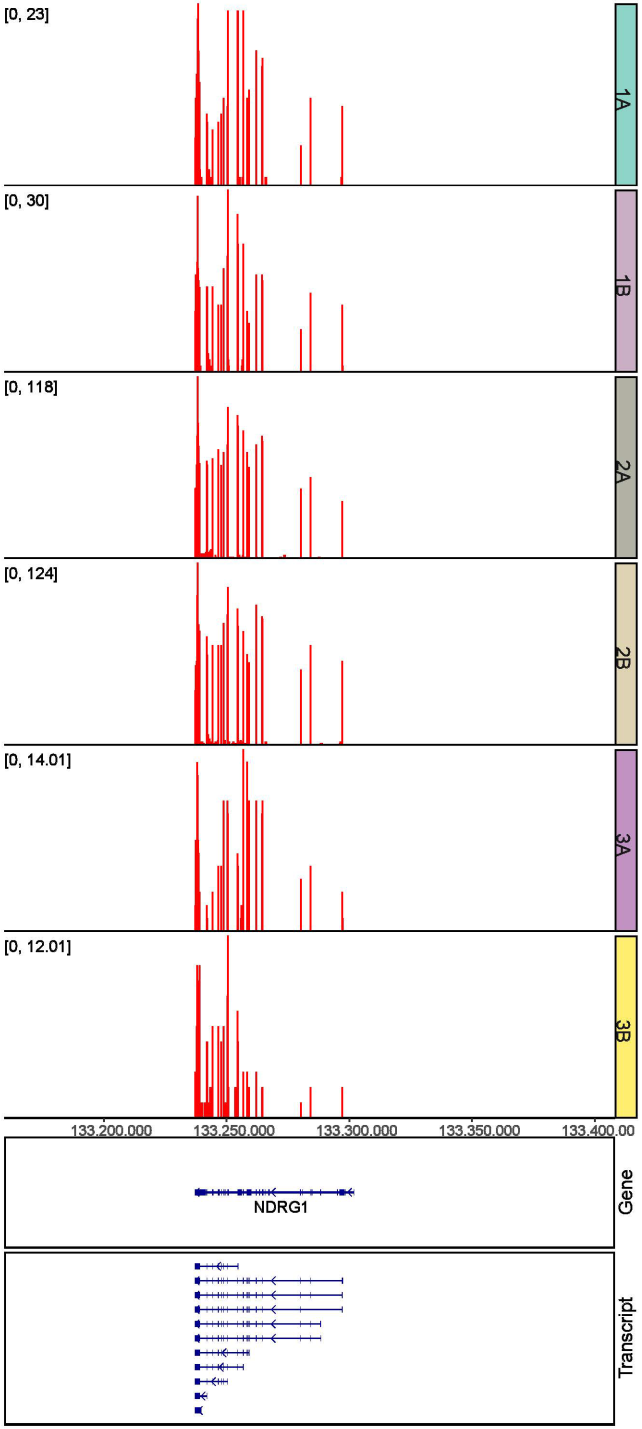


B

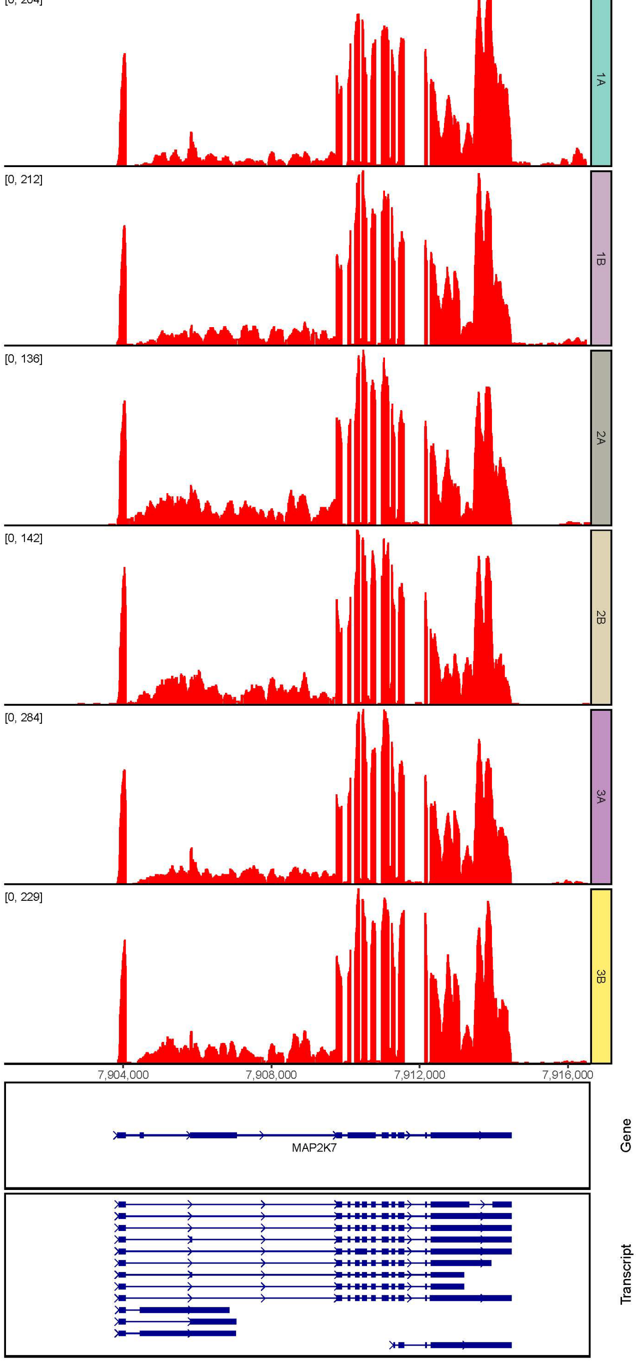


**Supplementary Figure 5**
